# LRRK2 kinase inhibition reverses G2019S mutation-dependent effects on tau pathology spread

**DOI:** 10.1101/2023.10.06.561190

**Authors:** Noah Lubben, Julia K. Brynildsen, Connor M. Webb, Howard L. Li, Cheryl E. G. Leyns, Lakshmi Changolkar, Bin Zhang, Emily S. Meymand, Mia O’Reilly, Zach Madaj, Daniella DeWeerd, Matthew J. Fell, Virginia M.Y. Lee, Dani S. Bassett, Michael X. Henderson

## Abstract

Mutations in leucine-rich repeat kinase 2 (LRRK2) are the most common cause of familial Parkinson’s disease (PD). These mutations elevate LRRK2 kinase activity, making LRRK2 kinase inhibitors an attractive therapeutic target. LRRK2 kinase activity has been consistently linked to specific cell signaling pathways, mostly related to organelle trafficking and homeostasis, but its relationship to PD pathogenesis has been more difficult to define. *LRRK2*-PD patients consistently present with loss of dopaminergic neurons in the substantia nigra but show variable development of Lewy body or tau tangle pathology. Animal models carrying LRRK2 mutations do not develop robust PD-related phenotypes spontaneously, hampering the assessment of LRRK2 inhibitors’ efficacy against disease processes. We hypothesized that mutations in LRRK2 may not be directly related to a single disease pathway, but instead may elevate susceptibility to multiple disease processes, depending on the disease trigger. To test this hypothesis, we have previously evaluated progression of α-synuclein and tau pathologies following injection of proteopathic seeds. We demonstrated that transgenic mice overexpressing mutant LRRK2 show alterations in the brain-wide progression of pathology, especially at older ages. Here, we assess tau pathology progression in relation to long-term LRRK2 kinase inhibition. Wildtype or LRRK2^G2019S^ knock-in mice were injected with tau fibrils and treated with control diet or diet containing LRRK2 kinase inhibitor MLi-2 targeting the IC50 or IC90 of LRRK2 for 3 to 6 months. Mice were evaluated for tau pathology by brain-wide quantitative pathology in 665 brain regions and subsequent linear diffusion modeling of progression. Consistent with our previous work, we found systemic alterations in the progression of tau pathology in LRRK2^G2019S^ mice that were most pronounced at 6 months. Importantly, LRRK2 kinase inhibition reversed these effects in LRRK2^G2019S^ mice, but had minimal effect in wildtype mice, suggesting that LRRK2 kinase inhibition is likely to reverse specific disease processes in G2019S mutation carriers, but additional work may be necessary to determine the potential effect in non-carriers. This work supports a protective role of LRRK2 kinase inhibition in G2019S carriers and provides a rational workflow for systematic evaluation of brain-wide phenotypes in therapeutic development.

## INTRODUCTION

Parkinson’s disease (PD) is a progressive disease characterized clinically by motor and non-motor symptoms and pathologically by the presence of Lewy bodies and Lewy neurites in the brain^1^. While most patients have no known genetic cause of disease, there are rare familial cases in which a genetic cause has been identified. Mutations in the leucine-rich repeat kinase 2 gene (*LRRK2*) are the most common cause of familial PD^2^. LRRK2 is a large protein with both kinase and GTPase domains and has been reported to participate in a number of cellular functions from lysosomal homeostasis to vesicular trafficking^3^. Most identified familial mutations, including the most common p.G2019S^4^, lead to elevated LRRK2 kinase activity^2^. For these reasons, LRRK2 kinase inhibitors have been pursued for the past 15 years as potential therapeutics for individuals with LRRK2 mutations^5^. One recent report has also suggested that LRRK2 kinase activity may be elevated in idiopathic PD patients^6^, extending the potential utility of LRRK2 kinase inhibitors to this much larger population. However, despite years of research, the connection between LRRK2 kinase activity and PD pathophysiology has remained unclear. Therefore, while several LRRK2 kinase inhibitors have shown excellent target engagement, it is uncertain what effect LRRK2 kinase inhibition may have on PD pathology or progression.

PD is characterized neuropathologically by the loss of dopaminergic neurons in the substantia nigra and the accumulation of α-synuclein aggregates in the form of Lewy bodies and Lewy neurites^7^. As disease progresses, up to 90% of PD patients also accumulate tau pathology, and tau pathology is associated with more extensive Lewy pathology and a worse prognosis for patients^8^. Since the identification of *LRRK2* mutations as a cause of familial PD, the pathology in *LRRK2*-PD has been less clear. Loss of substantia nigra neurons is ubiquitous in *LRRK2*-PD cases, but 21-54% of LRRK2 mutation carriers lack α-synuclein Lewy pathology^9–11^. Tau pathology is also common in *LRRK2*-PD cases, ranging from 79-100% ^11,12^, similar to the incidence of tau pathology in idiopathic PD, leading to the hypothesis that *LRRK2* mutations could predispose patients to the accumulation of either α-synuclein or tau pathologies. While multiple studies have suggested that *LRRK2* mutations are not independently associated with primary tauopathies such as progressive supranuclear palsy or corticobasal degeneration^13,14^, Alzheimer’s disease-type tau pathology still appears consistently *LRRK2*– and idiopathic PD^11^.

Recent work from several groups has therefore sought to determine the impact of LRRK2 kinase activity on α-synuclein and tau pathologies in animal models to provide insight into a relevant clinical outcome for LRRK2 kinase inhibition in clinical trials. Studies in mouse models of α-synucleinopathy have found that LRRK2^G2019S^ transgenic mice have similar overall pathology to wildtype mice, but with enhanced pathology in specific regions^15–17^. Similarly, in mouse models of tauopathy, LRRK2^G2019S^ appears to enhance tau pathology progression^18,19^. These studies have identified changes in pathology patterns that may be related to LRRK2 kinase activity and would therefore be amenable to reversal with LRRK2 kinase inhibition. However, a caveat of each of these studies was the use of transgenic overexpression of mutant LRRK2.

We therefore sought to assess the impact of LRRK2 kinase inhibition on tau pathology using a validated seed-based progressive tauopathy mouse model. Wildtype and LRRK2^G2019S^ knock-in (KI) mice were injected with Alzheimer’s disease (AD)-derived tau and concurrently treated with control diet or LRRK2 kinase inhibitor MLi-2 in diet targeting the IC50 or IC90 of LRRK2 based on previous research^20^. Mice tolerated chronic LRRK2 kinase inhibition well over 3-6 months dosing and MLi-2 was estimated to achieve 50-100% inhibition of LRRK2 kinase activity at medium and high doses, respectively. We then examined tau pathology systematically in 665 regions of the brain in all mice. LRRK2^G2019S^ mice showed a time-dependent enhancement of tau pathology, particularly in cortical brain regions, consistent with previous studies in transgenic mice. Remarkably, this alteration appeared kinase-dependent, since LRRK2 kinase inhibition was able to reverse this effect. We also found that LRRK2 kinase inhibition was not protective in wildtype mice. To gain further insight into the mechanism underlying these brain-wide changes, we computationally modeled pathology progression based on linear diffusion through the anatomical connectome. We found that diffusion through anatomical connectivity was a reliable predictor of pathology progression in all mice. LRRK2^G2019S^ mice had a reversal in spread direction induced by LRRK2 kinase inhibition. Together, the findings of this study demonstrate that LRRK2^G2019S^ has network-level effects on tau pathology progression that are reversible with LRRK2 kinase inhibition. Further, broad assessments of pathology and network modeling provide more systematic assessment of complex phenotypes in neurodegenerative disease models than more common targeted approaches.

## RESULTS

### Chronic LRRK2 inhibition is well-tolerated in wildtype and LRRK2^G2019S^ KI mice

We previously found that seeded tau pathology progresses through the mouse brain in a manner that is constrained by regional connectivity and vulnerability. In LRRK2^G2019S^ transgenic mice, pathology progresses even more in a retrograde direction^18^. To determine whether this is also true without overexpression of LRRK2 and whether LRRK2 kinase inhibition can reverse this effect, we injected LRRK2^G2019S^ knock-in (KI) mice^21^ and their wildtype littermates with tau paired helical filaments (PHFs) extracted from Alzheimer’s disease (AD) brains^22^ at 3 months of age (**Fig. 1A, Supplementary Table 1**). To assess the impact of LRRK2 kinase activity on observed effects, mice were given *ad libitum* access to diet formulated to deliver 10 or 60 mg/kg/day of the potent LRRK2 kinase inhibitor MLi-2^20^, or vehicle control diet one week prior to tau injection. Previous work showed that MLi-2 is fully brain penetrant^20^ and that these doses achieve approximately IC50 and full inhibition exposures in brain, respectively^23^. Mice were aged 3 or 6 months post-injection (MPI) and maintained on the altered diet for the duration of the study. Mice in all groups showed similar weight gain throughout the duration of the study (**Fig. 1B, 1C**). Mouse diet was weighed weekly, allowing an estimation of diet consumption and expected MLi-2 exposure. Based on these measures, mice fed with the 450 mg/kg MLi-2 diet maintained an exposure of 47-49 mg/kg/day (**Fig. 1D, 1E**), while mice fed with the 75 mg/kg MLi-2 diet averaged 8 mg/kg/day exposure (**Fig. 1E**).

**Figure 1.**
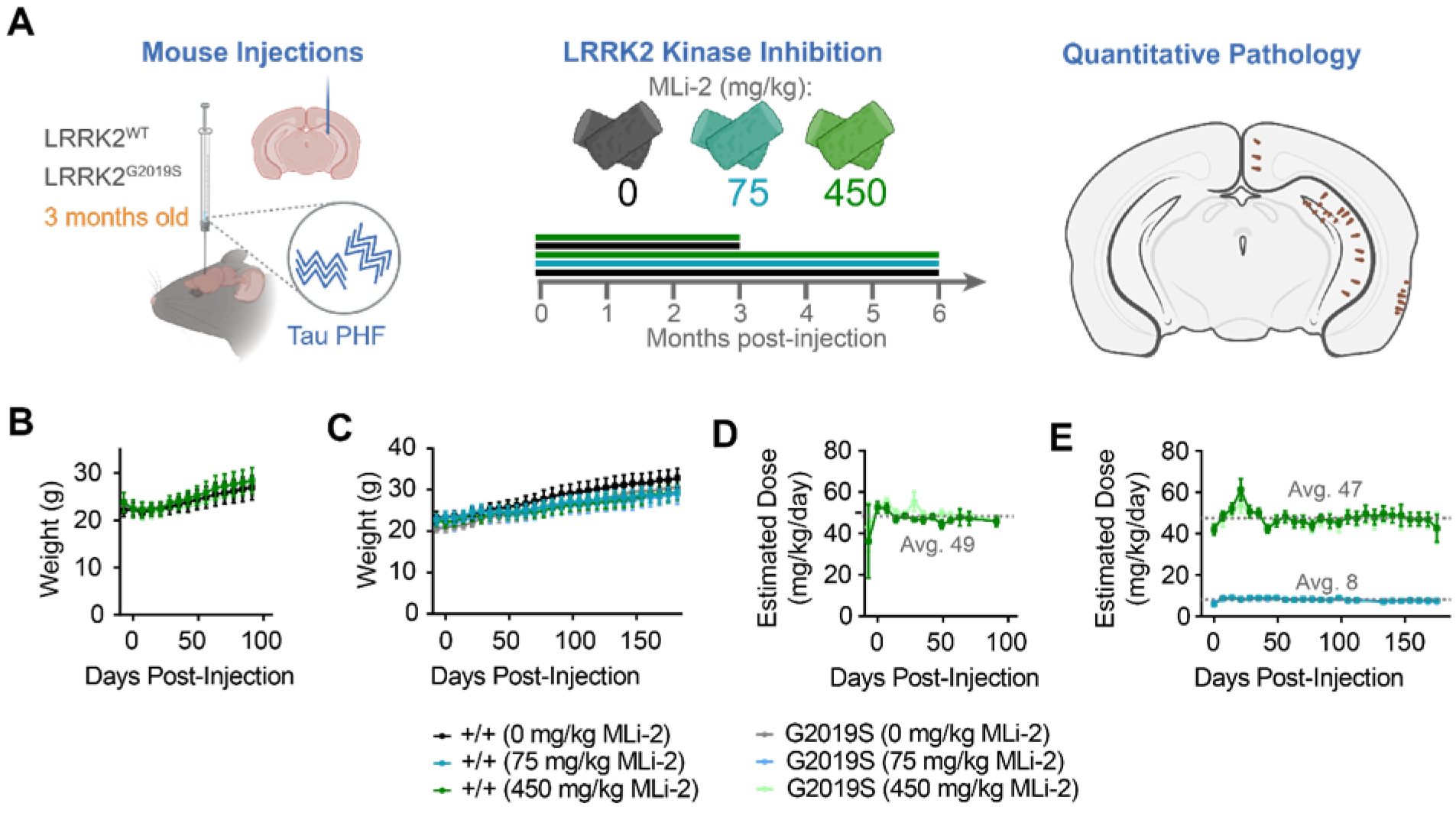
Chronic LRRK2 inhibition is well-tolerated in wildtype and LRRK2^G2019S^ KI mice. (**A**) LRRK2^G2Q19S^ KI mice and wildtype littermates were injected with tau paired helical filaments (PHFs) derived from Alzheimer’s disease brains at 3 months of age At the same time, mice were given access to diet with 0, 75, or 450 mg/kg MLi-2 incorporated Cohorts were aged 3 or 6 months as indicated Following aging, mice were sacrificed, and the primary endpoint of brain pathology was assayed Secondary studies of PK/PD and lung and kidney histology were also assayed (**B**) Average mice weights over 3 months of 0 or 450 mg/kg MLi-2 treatment (**C**) Average mice weights over 6 months of 0, 75, or 450 mg/kg MLi-2 treatment (**D**) Estimated exposure of mice in the 3-month cohorts to MLi-2 based on diet consumption (E) Estimated exposure of mice in the 6-month cohorts to MLi-2 based on diet consumption Mice on 450 mg/kg diet average 48 mg/kg/day exposure to MLi- 2, while mice on 75 mg/kg diet averaged 8 mg/kg/day exposure. Plots display the mean and standard error of each cohort at each timepoint.

### MLi-2 reduces total and pS935 LRRK2 and enlarges pneumocytes

To further explore the pharmacodynamic and pharmacokinetic profile of chronic MLi-2 exposure, plasma, kidney, and lung were collected from mice. LC-MS/MS analysis of plasma revealed terminal exposures of 8 nM and 23 nM unbound MLi-2 from mice fed 75 mg/kg and 450 mg/kg diet, respectively (**Fig. 2A**). Total LRRK2 and pS935 LRRK2 levels were assayed using an established MSD assay in the kidney as readouts of target engagement since the brain was fixed for histology. LRRK kinase inhibition significantly reduced LRRK2 levels, though not below the detection limit, whereas pS935 LRRK2 was dramatically reduced by MLi-2 treatment with pS935 LRRK2 becoming undetectable in mice treated with 450 mg/kg MLi-2 (**Fig. 2B, 2C**). Overall, mice on MLi-2 diet maintained partial to complete inhibition of LRRK2 kinase activity.

**Figure 2.**
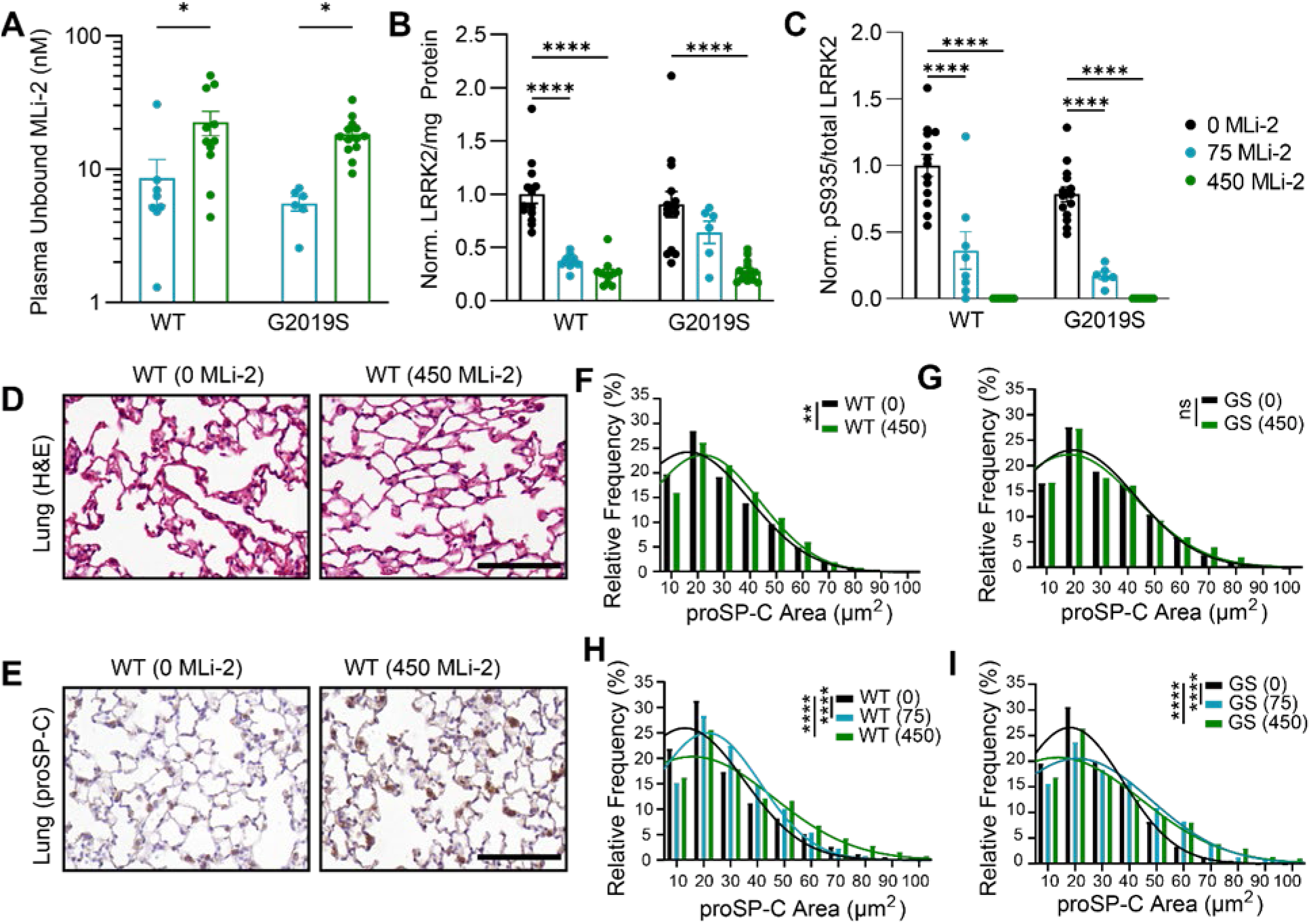
MLi-2 reduces total and pS935 LRRK2 and enlarges pneumocytes. (**A**) The pharmacokinetic profile of unbound MLi-2 was measured in the plasma of animals following in-diet dosing. *p<0.05, 2-Way ANOVA followed by Dunnetfs multiple comparison test to compare doses. (**B**) Total LRRK2 and (**C**) pS935 LRRK2 were reduced in the kidney following prolonged treatment with MLi-2. ****p<0.0001, 2-Way ANOVA followed by Dunnetfs multiple comparison test to compare doses. Animals treated for 3 months or 6 months were combined for these analyses. (**D**) Following hematoxylin and eosin staining, lung from mice treated with 450 mg/kg MLi-2 appear to have enlarged type II pneumocytes. Scale bar = 100 pm. (**E**) Pneumocytes were more easily observed and quantified following staining with an antibody targeting prosurfactant protein C (proSP-C). Scale bar = 100 pm. (**F**) proSP-C positive pneumocytes were detected and quantified using an automated algorithm. Size distributions of measured signal is plotted. Wildtype mice treated for 3 months showed a slight shift in distribution to larger sizes, (G) while G2019S mice did not show a significant shift. **p<0.01, Kolmogorov-Smirnov test. (H) After 6 months of treatment, wildtype and (**I**) G2019S mice showed larger distribution of proSP-C positive pneumocytes at both doses of MLi-2 compared to mice not exposed to MLi-2. ****p<0.0001, Kolmogorov-Smirnov test.

Several previous studies have shown that sustained LRRK2 kinase inhibition can lead to morphological phenotypes in the kidney and lung^20,24–26^. We therefore collected kidney and lung tissue and assayed each for morphological abnormalities. Kidney sections were stained with hematoxylin and eosin and showed no major morphological abnormalities (**Supplementary** Fig. 1). Lung tissue was stained with hematoxylin and eosin and also immunostained for prosurfactant protein C (proSP-C), a marker for pneumocytes. We found that mice treated with MLi-2, especially those treated with a high dose or for a longer time, showed enlarged pneumocytes (**Fig. 2D, 2E**). We performed a quantitative assessment of pneumocyte size based on proSP-C staining and found that the size distribution of pneumocytes in mice treated with MLi-2 was significantly increased in all groups, except for 3 MPI LRRK2^G2019S^ mice (**Fig. 2F-I**). These results are consistent with previous literature and suggest that a higher dose or more sustained LRRK2 inhibition will result in a stronger pneumocyte phenotype in the lung.

### Quantitative pathology to evaluate the development and spread of tau pathology

To investigate the effect of pharmacological inhibition of LRRK2 on the development and spread of tau pathology we used an established seed-based model of tauopathy that induces the misfolding of endogenous tau into hyperphosphorylated tau inclusions following injection of AD brain-derived tau into mice^22^. In the present study, we injected LRRK2^G2019S^ knock-in and WT littermates with AD brain-derived PHFs and then aged them to either 3 or 6 months MPI. AD PHF tau induces misfolding of endogenous mouse tau and subsequent progression of phosphorylated tau pathology throughout the mouse brain^18,22^. Brains were sectioned, and 12 representative sections were selected and stained for phosphorylated tau pathology (AT8, pS202/T205 tau). To quantify phosphorylated tau pathology throughout the brain, we implemented a segmentation and brain registration platform adapted from the QUINT workflow^27^ (**Supplementary** Fig. 2). Segmentation was performed in QuPath using an optical density pixel threshold. In parallel, tissue sections were placed in a three-dimensional atlas using QuickNII^28^, and nonlinear warp transformations were performed in VisuAlign. Segmentations were aligned to registered anatomical regions and quantified in Nutil^29^ software. This strategy enabled us to produce highly-reproducible measure of tau pathology burden in 665 anatomical regions across the brain (**Supplementary** Fig. 3).

### LRRK2^G2019S^ causes time-dependent alterations in cortical tau pathology

We have previously demonstrated that LRRK2^G2019S^ overexpressing mice have alterations in tau pathology that are most pronounced at later timepoints^18^. Since it was not feasible in the current study design to include 9 MPI, we assessed genotype-related differences at 3 and 6 MPI using semi-parametric ordinal regressions, while accounting for multiple comparisons. We first assessed tau pathology differences by sex in wildtype mice (**Supplementary** Fig. 4). Interestingly, we found increases in subcortical pathology at 3 MPI, and less pronounced increases at 6 MPI in male mice. Therefore, all ordinal regression also accounted for sex in estimating differences. We found minimal difference between wildtype and LRRK2^G2019S^ mice at 3 MPI (**Supplementary** Fig. 5**, 6**). However, by 6 MPI, pathology was significantly higher in many cortical regions of LRRK2^G2019S^ mice, with the exception of entorhinal cortex (**Fig. 3A, 3B**). At this timepoint, pathology has begun to progress past the initial injection sites and their directly connected areas (hippocampus, entorhinal cortex, supramammillary nucleus), and the increase in pathology in LRRK2^G2019S^ mice is observed in more distally connected sites. Based on previous data in LRRK2^G2019S^ transgenic mice, we would expect the two genotypes to further differentiate at later timepoints, consistent with a role for LRRK2 in pathology progression, but not in pathology initiation.

**Figure 3.**
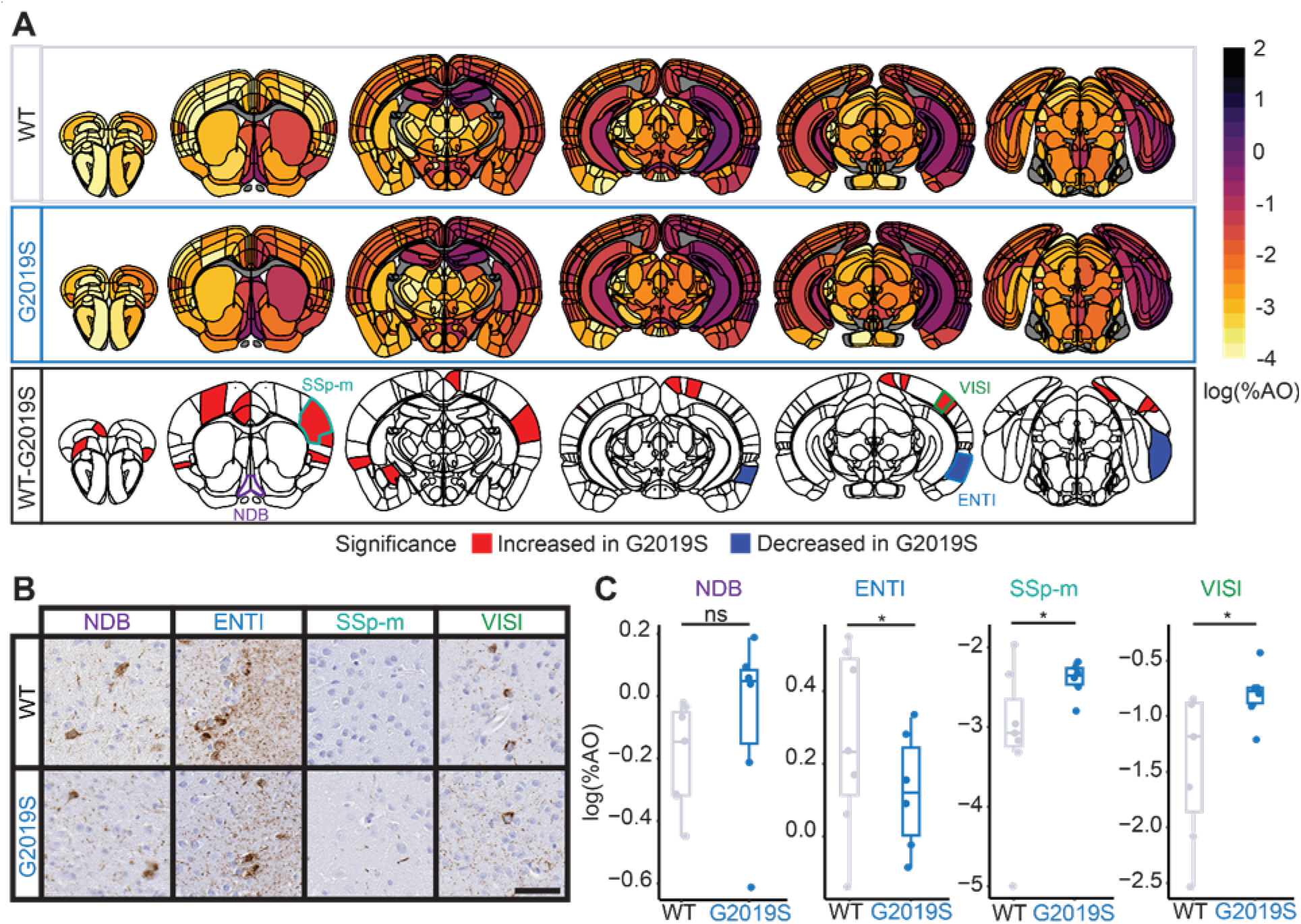
LRRK2^G2019S^ mice show time-dependent alterations in cortical pathology. (**A**) Anatomic heatmaps of mean regional tau pathology (ATS, pS202/T205 tau) shown as log (% area occupied) at 6 MPI and second-generation p-vaiues, calculated using ranged robust linear regression, of regional statistical significance of wildtype mice compared to G2019S mice (6p=0). Tau pathology was not quantified in white matter regions, so they are plotted as gray Analyzed regions shown below are outlined. NDB: diagonal band nucleus, ENTI: entorhinal area, lateral part, SSp-m: primary somatosensory area, mouth, ViSI: lateral visual area. (**B**) Representative images of selected regional tau pathology. Scale bar = 50 pm. (**C**) Quantification of selected brain regions. Regional tau shown as log (% area occupied) ns = not significant (δp≠0), *: significant (δp=0) n= 7 (WT) and 6 (G2019S)

### LRRK2 kinase inhibition shows no protective effect in wildtype mice

If LRRK2 kinase activity is implicated in cases lacking a LRRK2 mutation, then LRRK2 kinase inhibition may have a beneficial effect on pathology in wildtype mice. To test this hypothesis, we next compared wildtype mice treated with LRRK2 kinase inhibitors to controls. Again, we found minimal differences in tau pathology at 3 MPI (**Supplementary** Fig. 7). However, at 6 MPI, we observed a broad elevation in tau pathology in the 75 mg/kg dose group (**Fig. 4A, Supplementary** Fig. 8). Histologically, this elevation appeared to be primarily driven by neuritic pathology in those regions (**Fig. 4B**). Remarkably, this effect was nearly absent in the 450 mg/kg dose group, so does not appear to be a dose-dependent phenotype. At either timepoint, these data do not support a protective role for LRRK2 kinase inhibitors in a wildtype mouse of with tau pathology.

**Figure 4.**
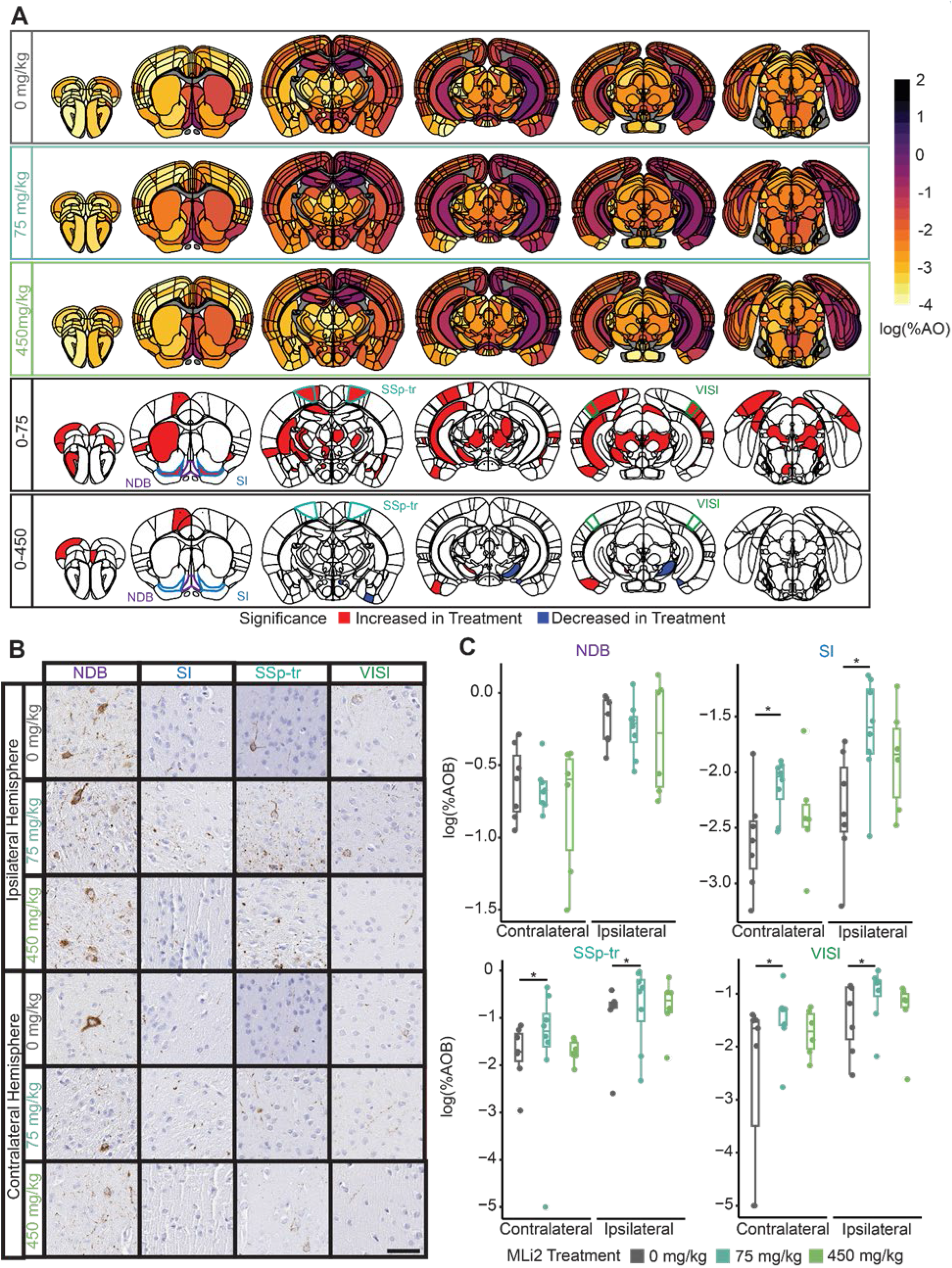
LRRK2 kinase inhibition shows no protective effect in wildtype mice. (**A**) Anatomic heatmaps of mean regional tau pathology shown as log (% area occupied) at 6 MPI and second-generation p-values of regional statistical significance, calculated using ranged robust linear regression, of WT mice with 0 mg/kg MLi-2 compared to wildtype mice treated with 75 or 450 mg/kg MLi-2 (5p=0). Tau pathology was not quantified in white matter regions, so they are plotted as gray. Analyzed regions shown below are outlined. NDB: diagonal band nucleus, SI: substantia innominata, SSp-tr: primary somatosensory area, trunk, VISI: lateral visual area. (**B**) Representative images of selected regional tau pathology. Scale bar = 50 pm. (C) Quantification of selected brain regions. Regional tau shown as log (% area occupied) ns = not significant (δp≠0), *: significant (δp=0) n= 7 (0 mg/kg MLi-2), 8 (75 mg/kg MLi-2), or 6 (450 mg/kg MLi-2)

### LRRK2 kinase inhibition reduces cortical pathology in LRRK2^G2019S^ mice

LRRK2^G2019S^ and related mutations have been demonstrated to elevate LRRK2 kinase activity. Therefore, we next assessed the ability for LRRK2 kinase inhibitor MLi-2 to alter tau pathology progression in LRRK2^G2019S^ knock-in animals. Similar to the genotype effect, we saw minimal treatment effect at 3 MPI (**Supplementary** Fig. 7), suggesting that kinase activity is not essential for initial tau seeding. However, at 6 MPI, there was a broad decrease in cortical tau pathology in mice treated with 450 mg/kg MLi-2, but not 75 mg/kg MLi-2 (**Fig. 5A, 5B**). The main exception to this pattern was the entorhinal cortex, which is highly connected to the injection site. It is interesting that full inhibition of LRRK2 kinase activity seems to be required for this effect, despite the fact that 75 mg/kg MLi-2 should have reduced LRRK2 kinase activity below wildtype levels (**Fig. 1C**). The increase in the entorhinal area and decrease in other cortical regions suggests that primary pathology seeding is not affected by LRRK2 kinase activity, unlike the progression to other brain regions. However, these broad changes can be difficult to assess region-by-region, so we turned to computational modeling of progression.

**Figure 5.**
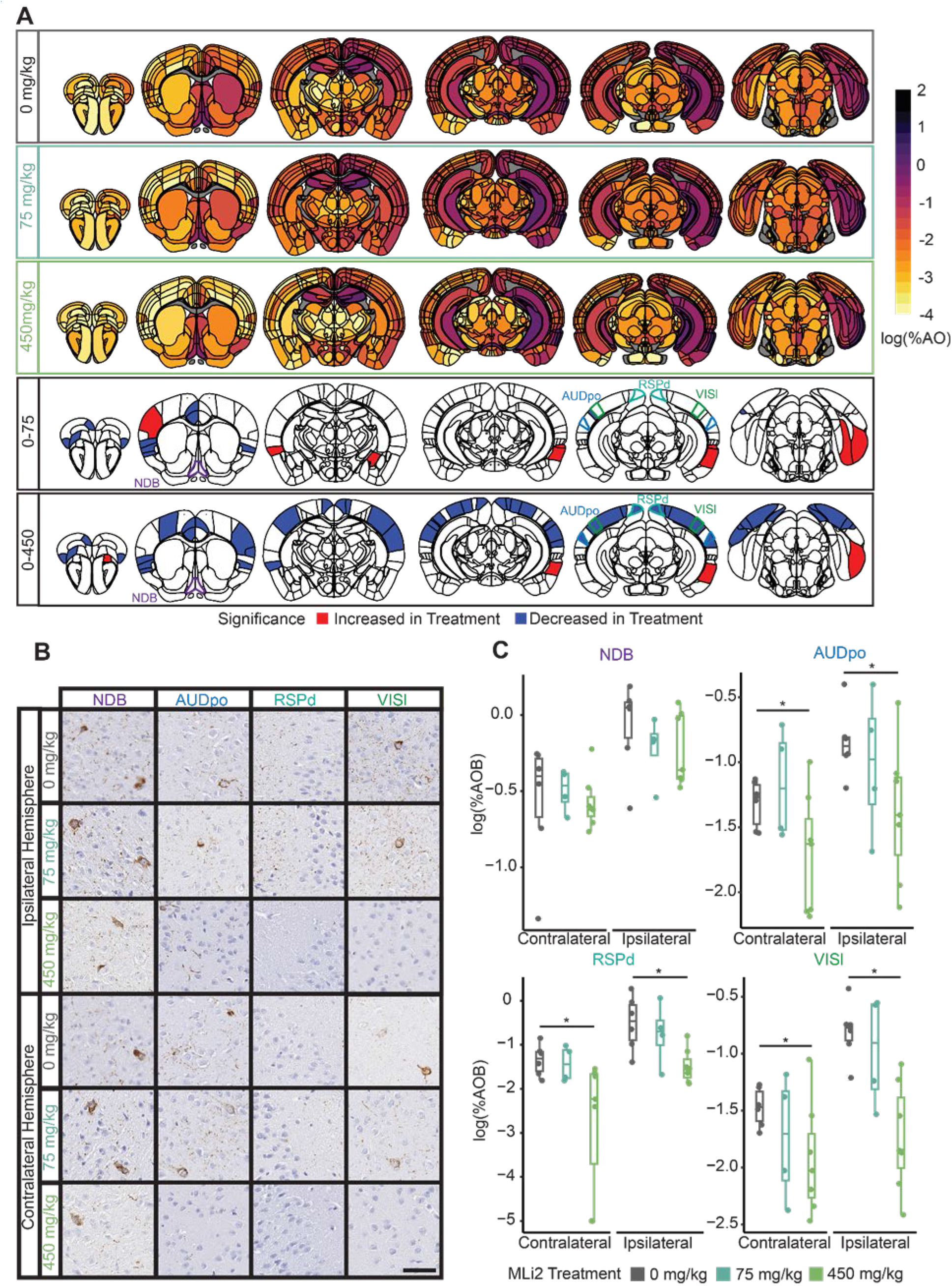
LRRK2 kinase inhibition reduces cortical pathology in LRRK2^G2019S^ mice. (**A**) Anatomic heatmaps of mean regional tau pathology shown as log (% area occupied) at 6 MPI and second-generation p-values, calculated using ranked robust linear regression, of regional statistical significance of G2019S mice with 0 mg/kg MLi-2 compared to G2019S mice treated with 75 or 450 mg/kg MLi-2 (6p=0). Tau pathology was not quantified in white matter regions, so they are plotted as gray. Analyzed regions shown below are outlined. NDB: diagonal band nucleus, AUDpo: posterior auditory area, RSPd: retrosplenial area, dorsal part, VISI: lateral visual area. (**B**) Representative images of selected regional tau pathology. Scale bar = 50 pm. (C) Quantification of selected brain regions. Regional tau shown as log (% area occupied) ns = not significant (δp≠0), *: significant (δp=0) n= 6 (0 mg/kg MLi-2), 4 (75 mg/kg MLi-2), or 7 (450 mg/kg MLi-2).

### LRRK2^G019S^ and LRRK2 kinase inhibition impacts network progression of tau pathology

Previous work using computational modeling of intracellular pathology progression has demonstrated that anatomical connectivity is a major constraint on pathology progression^16,18,30^. While this type of modeling is useful to understand how pathology progresses through the brain and factors underlying regional vulnerability, we hypothesized that it could also be useful in understanding the impact of compound administration on network dynamics of pathology spread. We therefore implemented linear diffusion modeling on the brain-wide tau pathology data collected in this study.

Since this was the first collection of high resolution pathology data (665 regions versus our 134 regions previously^18^), we first confirmed that a linear diffusion model with anatomical connectivity as an input layer would be able to predict pathology in each of the genotypes and treatment groups. Notably, only 420 of the 665 regions we measured had corresponding connectivity data, largely due to the lack of cortical laminar connectivity data. Consistent with our previous study, we determined that our bidirectional spread model strongly predicted levels of tau pathology at 3 and 6 MPI in both WT and LRRK2^G2019S^ mice in all treatment groups (**Fig. 6A**, **Supplementary** Fig. 9). Overall model fit appeared to be unaffected by treatment with MLi-2. We validated the specificity of our model by comparing fits obtained with the actual seed regions to those obtained by randomly selecting 500 alternate sets of 5 seed regions. We found that using the true seed regions yielded a significantly better fit than using random seeds (*p* < 0.002; **Fig. 6B**), indicating that the model’s performance was specific to the experimental seed sites. Finally, we compared model fits obtained from our bidirectional spread model to those obtained by modeling spread only in the anterograde direction, only in the retrograde direction, or based on Euclidean distance. The bidirectional spread model yielded a significantly better fit than models based on anterograde spread or Euclidean distance at 3 MPI, and its performance was superior to anterograde, retrograde, and distance-based models at 6 MPI (*p* < 0.002).

**Figure 6.**
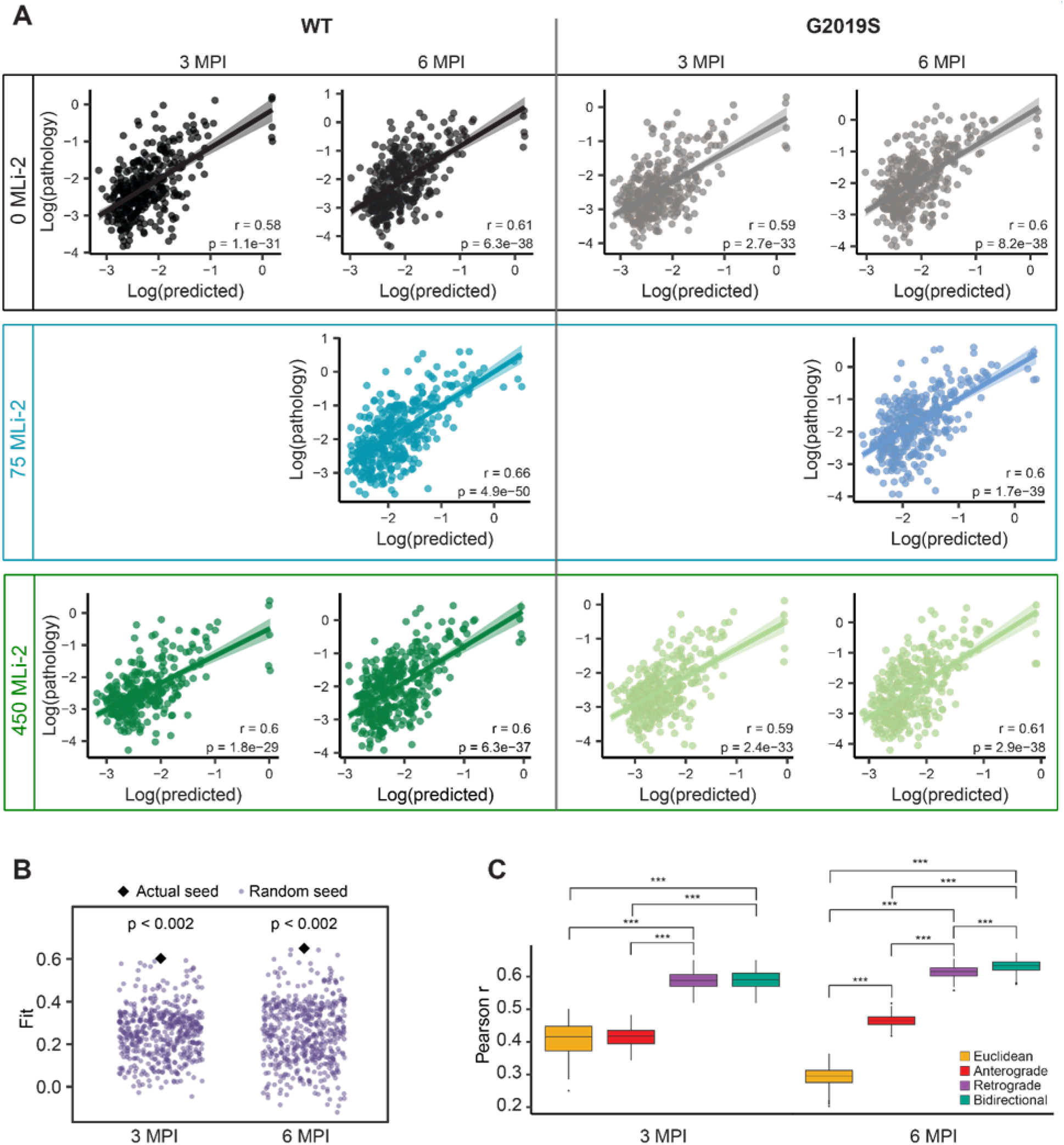
Modeling tau pathology spread. (**A**) Predictions of log tau pathology from linear diffusion models based on bidirectional (anterograde and retrograde) anatomical connections. Solid lines represent the line of best fit, and shading represents 95% confidence intervals. (**B**) Comparison of Pearson’s r values obtained by fitting bidirectional spread models using actual (black diamond) and alternate (purple points) seed regions. Using the true seeds yielded a better fit than using random seeds at both 3 and 6 MPI. (**C**) Comparison of Euclidean, anterograde, retrograde, and bidirectional model fits in 500 held-out samples. Retrograde and bidirectional spread models performed significantly better than the anterograde and Euclidean spread models at 3 MPI, and the bidirectional model performed significantly better than all other models at 6 MPI (***p< 0.002).

To assess the impact of MLi-2 treatment on mechanisms of pathology spread, we fit the bidirectional spread model on bootstrapped samples of data from mice in each treatment group to obtain distributions of Pearson’s *r*, the diffusion rate constant, and standardized regression weights (**Fig. 7A, 7B**). We observed no difference in the overall fit of the bidirectional diffusion model across genotypes and treatment groups and no difference in the diffusion rate constant. LRRK2 kinase inhibition in wildtype mice had subtle effects on standardized β weights that were not significant. However, retrograde β weights were lower in LRRK2^G2019S^ mice, and MLi-2 treatment elevated those weights to levels more comparable to wildtype mice.

**Figure 7.**
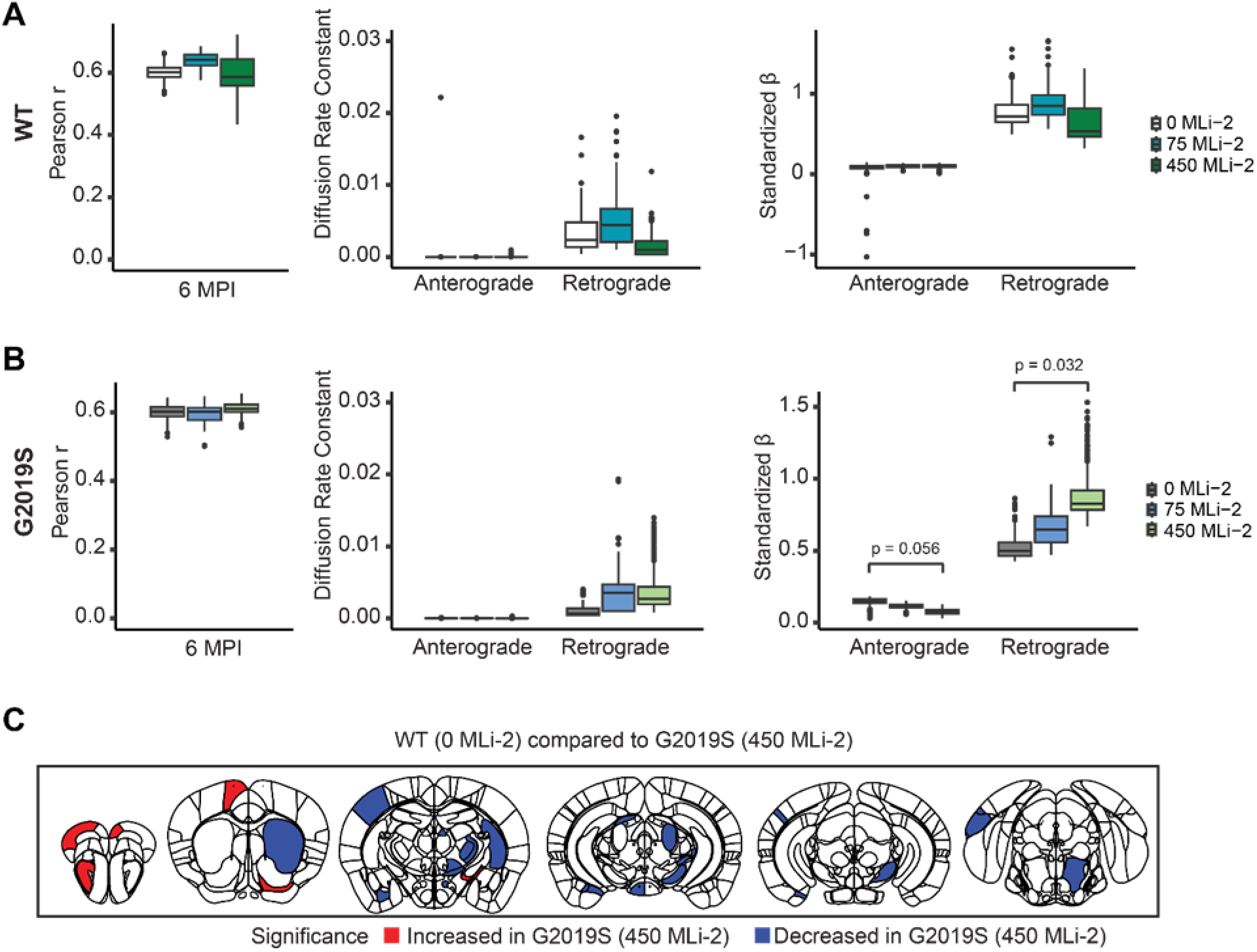
LRRK2^G019S^ and LRRK2 kinase inhibition impacts network progression of tau pathology. Comparisons of Pearson r, diffusion rate constant and standardized p in WT mice (**A**) treated with 0, 75, or 450 mg/kg MLi-2 and LRRK2^G2O19S^ mice (**B**) treated with 0, or 450 mg/kg MLi-2. Distributions were generated for each group by obtaining 500 bootstrapped resamples and fitting the bidirectional diffusion model for each resample. Since data for the 75 mg/kg dose of MLi-2 were only available at 6 MPI, bootstrapping for each group was performed using only data from this time point. (C) Results of ranked robust linear regression analysis of regional tau pathology in WT mice under control treatment (0 mg/kg MLi-2) compared to LRRK2^G2O19S^ mice treated with 450 mg/kg MLi-2. Blue and red colored regions have second generation p-values equal to 0.

Modeling results support a shift in progression parameters induced in LRRK2^G2019S^ animals that is reversed by LRRK2 kinase inhibition. To assess this shift in individual regions, we compared wildtype mice on control diet to LRRK2^G2019S^ mice treated with 450 mg/kg MLi-2 for 6 months (**Fig. 7C**). Tau pathology was almost completely normalized by MLi-2 treatment, with almost all regions showing either no difference from wildtype mice or a significant reduction in pathology. Together our data support a role for LRRK2 kinase activity in the progression of tau pathology and show that these effects can be reversed by chronic LRRK2 kinase inhibition, demonstrating the value of broad pathology evaluation and computational modeling in the evaluation of therapeutics (**Fig. 8**).

**Figure 8.**
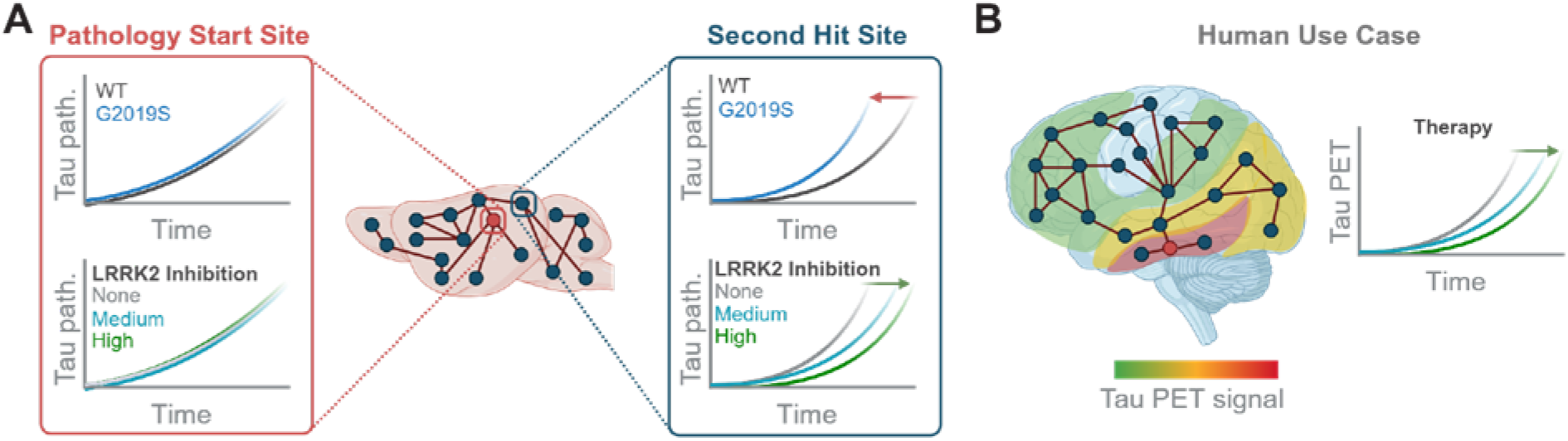
Model of LRRK2 impact on pathology progression. (**A**) Graphical summary of major study findings. Near the pathology start site and highly connected regions, there was minimal effect of either LRRK2^G2O19S^ expression or LRRK2 kinase inhibition. However, in other regions, primarily cortical regions, there was an acceleration of tau pathology in LRRK2^G2O19S^ mice that was reversed with LRRK2 kinase inhibition. (**B**) Proposed use of regional measurements and progression modeling to assess the impact of therapies either preclinically or clinically. Tau pathology can be assessed in living patients by PET imaging. Further, tau PET signal progresses in a predictable fashion, with individual variability. We propose that rather than assessing single regions, it may be more valuable to sample broadly and model the impact of therapies on progression.

## DISCUSSION

LRRK2 mutations are the most common cause of familial PD, and the development of LRRK2 kinase inhibitors has been a major focus in PD research for the past 15 years^31^. Yet, the field has been slowed by two major roadblocks—concerns over the safety of LRRK2 kinase inhibition, and the lack of a validated biological readout for compound efficacy. Several studies have sought to understand the safety of LRRK2 inhibition^32^, including recent Phase I clinical trials that assessed multiple doses in humans^33^. These safety studies concluded that while there are reproducible histological phenotypes associated with LRRK2 kinase inhibition, it seems possible to wash out these effects, and there is no overt change in lung function in animal studies or humans. However, the role of LRRK2 in PD pathophysiology has remained elusive, making testing of compounds on disease measures difficult. In the current study, we attempted to assess the impact of LRRK2^G2019S^ and LRRK2 kinase inhibition on tau pathology progression. We find that a biological effect was only observable when considering a large number of brain regions at extended timepoints, suggesting that it may be beneficial to expand compound efficacy studies outside of individual brain regions or timepoints.

We have previously demonstrated an effect of LRRK2^G2019S^ overexpression on the progression of tau pathology in AD PHF injected mice^18^. Notably, effects on pathology were only apparent at later timepoints. This absence of early change is consistent with a recent study that injected α-synuclein PFFs in the hippocampus of LRRK2 knockout and LRRK^G2019S^ knock-in mice, and observed no differences in α-synuclein pathology at 1 MPI^34^. Importantly, we and others often measure pathology in the injected region or regions with a high number of projections to the injected region. However, in the current and other studies^18,19^, these regions show minimal changes in pathology. Together these data suggest that LRRK2 may enhance tau progression to distal sites, while reducing pathology in primary hit sites (entorhinal cortex). The cellular mechanism for this network-level phenomenon remains unclear, but could be related to increased synaptic vesicle release^35^ or altered autophagosome trafficking^36^ that has been observed in LRRK2^G2019S^ neurons. Importantly, LRRK2 kinase inhibition was able to reverse pathology changes in LRRK2^G2019S^ mice but showed no beneficial effect in wildtype mice. While preclinical studies should be extrapolated with caution, our study suggests that LRRK2 kinase inhibitors may be beneficial in LRRK2^G2019S^ carriers, but not in non-carriers. However, there are many important differences between human PD and mouse models that could yield different outcomes in patients than in mice.

In addition to broad pathology assessment, we also employed computational modeling of pathology progression to understand the impact of LRRK2^G2019S^ and kinase inhibition in our animal model. We implemented an improved pathology quantitation workflow to increase brain coverage in this study to 665 regions. Given the novelty of this approach, we first wanted to validate its ability to predict pathology progression based on anatomical connectivity. We show a similar ability as our previous studies to predict pathology patterns, but with improved statistical confidence. Network modeling indicated that there were differences in standardized β weights, or the overall directionality of pathology progression, in LRRK2^G2019S^ mice that were reversed by LRRK2 kinase inhibition. Yet, this model was based only on the strength of inter-regional connections. Future models incorporating gene expression or functional activity may improve predictivity^37^ and thereby improve our understanding of the effects of gene mutations and therapies on pathology progression.

Together, broad assessments of pathology and network modeling can provide novel insights into disease progression and the impact of therapies^37^. In addition to their utility in preclinical models, we argue that they may prove useful in clinical evaluation of therapies, although further development in this area would be required. Some therapies would be expected to act throughout the brain and would make broad assessments unnecessary. For example, anti-amyloid β immunotherapies can clear amyloid β plaques throughout the brain^38^. However, other therapies may reduce progression of pathology, while not impacting pathology in baseline regions. Tau pathology is measurable in living patients by positron emission tomography (PET) and may prove a valuable biomarker of disease progression^39,40^. One can imagine a scenario in which a therapy does not change pathology at primary sites observed with a baseline PET, but subsequent PET imaging shows a reduction in non-primary regions compared to placebo. It is unclear if tau pathology is substantial enough in PD patients, or whether tau PET is sensitive enough to differentiate more subtle pathology changes, but this type of secondary analysis may be broadly applicable in neurodegenerative disease, if not in living patients, then possibly in post-mortem neuropathological studies.

There are several limitations to the current study. To make this study feasible within time constraints, we only measured two timepoints with a single LRRK2 kinase inhibitor. Longer timepoints may be necessary to provide a full picture of the impact of LRRK2 on pathology progression, and it would be important to show biological efficacy for compounds that are bound for the clinic. We also only use one variable, pS202/T205 tau (AT8), as a pathology measure in this study. The amount of time it takes to segment and register brain sections was prohibitive to assessing more than one measure, but improvements in this process may make this technique more streamlined in the future. We started LRRK2 kinase inhibition concurrent with tau pathology induction in this study, so the results could be perceived as preventative, not protective. However, given that minimal effects were seen at 3 MPI, we hypothesize that treatment could be delayed for some time with similar results.

In conclusion, we found that the LRRK2^G2019S^ mutation accelerates tau pathology progression, which was reversed by LRRK2 kinase inhibition. We also establish a novel workflow for the assessment of potential therapies on the network-level that we believe will be useful for future research.

## GENERAL

We would like to thank the patients and families who donate tissue for this research, without whom this study would not have been possible. We would like to thank members of the laboratory for their feedback in developing this manuscript and the Van Andel Institute Pathology and Biorepository Core (RRID:SCR_022912) for their assistance with tissue sectioning. Several images were created with BioRender.com.

## AUTHOR CONTRIBUTIONS

Conceptualization, M.X.H..; Methodology, N.L., J.K.B., C.E.G.L., L.C., B.Z., M.X.H.; Software, J.K.B., Z.M., D.D., D.S.B.; Formal Analysis, N.L., J.K.B., H.L.L., C.E.G.L., Z.M., M.X.H.,; Investigation, N.L., J.K.B., C.W., C.E.G.L., L.C., B.Z., E.S.M., M.O., M.X.H.; Resources, M.J.F., V.M.Y.L., D.S.B., M.X.H.; Writing-Original Draft, N.L., J.K.B., M.X.H.; Writing-Review and Editing, All; Visualization, N.L., J.K.B., M.X.H.; Supervision, M.J.F., V.M.Y.L., D.S.B., M.X.H.; Funding Acquisition, M.J.F., V.M.Y.L., D.S.B., M.X.H.

## DATA AVAILABILITY

The data that support the findings of this study are available together with the code used for data analysis here: https://github.com/jkbrynildsen/tau-spread.

## CODE AVAILABILITY

Primary code used to analyze data and generate linear diffusion models is available here: Initial conversion of Nutil files to matrix output and grid heatmap generation: https://github.com/DaniellaDeWeerd/NutilToUsable. Brain heatmaps and differential analysis of area occupied: https://github.com/vari-bbc/Mouse_Brain_Heatmap. Computational models: https://github.com/jkbrynildsen/tau-spread.

## MATERIALS AND METHODS

### Mice

All housing, breeding, and procedures were performed according to the NIH Guide for the Care and Use of Experimental Animals and approved by the University of Pennsylvania’s Institutional Animal Care and Use Committee. LRRK2^G2019S^ knock-in mice (C57BL/6-*Lrrk2^tm^*^4^*^.1Arte^*, RRID:IMSR_TAC:13940) have been previously described^21^. Mice were bred as heterozygotes, and homozygous (+/+ or G2019S/G2019S) littermates were used for experiments. Both male (n=30) and female (n=35) mice were used and were 3-4 months old at the time of injection.

### MLi-2 Compound Administration

MLi-2 was synthesized at Merck Research Laboratories (Boston, MA). MLi-2 was dosed in rodent chow, prepared at Research Diets (New Brunswick, NJ), and formulated to include 0, 75, or 450 mg/kg MLi-2. The base for each diet was the vehicle diet (D01060501; Research Diets). Achieved dose was estimated based on weekly diet consumption per mouse and mouse weight.

MLi-2 was synthesized at Merck & Co., Inc. (Boston, MA, USA). Diet was prepared at Research Diets (New Brunswick, NJ), and formulated to deliver 0, 10, or 60 mg/kg/day MLi-2, based on an average daily food consumption of 4g per 30g mouse. The base for each diet was the vehicle diet (D01060501; Research Diets).

### Human Tissue

All procedures were done in accordance with local institutional review board guidelines of the University of Pennsylvania. Written informed consent for autopsy and analysis of tissue sample data was obtained either from patients themselves or their next of kin. All cases used for extraction of PHF tau (**Supplementary Table 1**) were Braak stage VI and were selected based upon a high burden of tau pathology by immunohistochemical staining. Cases used for extraction were balanced by sex (female = 2; male = 1) and were frozen an average of 14 hours post-mortem. After extracting all three cases and confirming the seeding capacity of tau, cases were pooled for injection and all mice were injected with the same AD PHF pool.

### Human Brain Sequential Detergent Fractionation

Frozen postmortem human frontal or temporal cortex brain tissue containing abundant tau-positive inclusions was selected for sequential extraction based on IHC examination of these samples as described^41^ using previously established methods. These brains were sequentially extracted with increasing detergent strength as previously described ^22^. After thawing, meninges were removed and gray matter was carefully separated from white matter. Gray matter was weighed and suspended in nine volumes (w/v) high salt (HS) buffer (10 mM Tris-HCL (pH 7.4), 800 mM NaCl, 1 mM EDTA, 2 mM dithiothreitol [DTT], protease and phosphatase inhibitors and PMSF) with 0.1% sarkosyl and 10% sucrose, followed by homogenization with a dounce homogenizer and centrifugation at 10,000 x *g* for 10 minutes at 4°C. The resulting pellet was re-extracted with the same buffer conditions and the supernatants from all extractions were filtered and pooled.

Additional sarkosyl was added to the pooled supernatant to reach a final concentration of 1% and the supernatant was nutated for 1 hour at room temperature. The samples were then centrifuged at 300,000 x *g* for 60 minutes at 4°C. The pellet, which contains pathological tau, was washed once with PBS and resuspended in 100 μL of PBS per gram of gray matter by passing through a 27G/0.5 inch needle. The pellets were further suspended by brief sonication (QSonica Microson™ XL-2000; 20 pulses; setting 2; 0.5 sec/pulse). The suspension was centrifuged at 100,000 x *g* for 30 minutes at 4°C. The pellet was suspended in one-fifth to one-half the pre-centrifugation volume, sonicated briefly (60-120 pulses; setting 2; 0.5 sec/pulse) and centrifuged at 10,000 x *g* for 30 minutes at 4°C. The final supernatant was utilized for all studies and is referred to as AD PHF tau. All extractions were characterized by Western blotting for tau, sandwich ELISA α-synuclein and Aβ 1–40, Aβ 1–42, and validated by immunocytochemistry in primary neurons from non-transgenic mice. For the extractions used in this study, tau constituted 9-11% of the total protein, while α-synuclein and Aβ constituted 0.008% or less of total protein.

### Sandwich ELISA

Characterization of α-synuclein and Aβ 1–40, Aβ 1–42 from AD PHF preparations by sandwich ELISA has been previously described^22^. Assays were performed on 384-well MaxiSorp clear plates (ThermoFisher). Plates were coated with well-characterized capture antibodies (α-synuclein: Syn9027; Aβ 1–40/ Aβ 1–42: Ban50) at 4°C overnight, washed, and blocked with Block Ace (AbD Serotec) overnight at 4°C. AD PHF preparations were diluted at 1:100 and added to plates alongside serial dilutions of recombinant α-synuclein, recombinant T40, or peptides for Aβ 1-40 and 1-42 monomeric standards. Plates were incubated overnight at 4°C, then washed and incubated with reporter antibodies (tau: BT2+BT7; α-synuclein: MJF-R1; Aβ 1–40: BA27; Aβ 1–42: BC05) overnight at 4°C. Plates were washed and incubated with HRP-conjugated secondary antibodies for 1 hour at 37°C. Plates were developed with 1-Step Ultra TMB-ELISA Substrate Solution (Thermo Fisher Scientific) for 10-15 minutes. The reaction was quenched with 10% phosphoric acid and read at 450 nm on a plate reader (M5, SpectraMax).

### Stereotaxic Injection

All surgery experiments were performed in accordance with protocols approved by the Institutional Animal Care and Use Committee (IACUC) of the University of Pennsylvania. AD PHF tau from individual extractions was vortexed and diluted with DPBS to 0.4 mg/mL. Tau was sonicated in a cooled bath sonicator at 9°C (Diagenode Bioruptor®; 20 cycles; setting medium; 30 seconds on, 30 seconds off). Mice were injected when 3-4 months old. Mice were deeply anaesthetized with ketamine/ xylazine/ acepromazine and injected unilaterally by insertion of a single needle into the right forebrain (coordinates: –2.5 mm relative to Bregma, +2.0 mm from midline) targeting the hippocampus (2.4 mm beneath the skull) with 1 µg tau (2.5 µL). The needle was then retracted to 1.4 mm beneath the skull, targeting the overlaying cortex and another 1 µg tau (2.5 µL) was injected. The needle was left in place for 2 minutes following the injection. Injections were performed using a 25 µL syringe (Hamilton, NV) at a rate of 0.4 µL/minute.

### Hematoxylin and Eosin Staining

Mice were perfused transcardially with PBS. Lung and kidney tissues were removed from mice and post-fixed in 10% neutral-buffered formalin (NBF). Lung tissues were first inflated by injection with 10% NBF. After perfusion and fixation, both types of tissues were processed into paraffin via sequential dehydration and perfusion with paraffin under vacuum (70% ethanol for 2 hours, 80% ethanol for 1 hour, 95% ethanol for 1 hour, 95% ethanol for 2 hours, 3 times 100% ethanol for 2 hours, xylene for 30 minutes, xylene for 1 hour, xylene for 1.5 hours, 3 times paraffin for 1 hour at 60°C). Following fixation and paraffin processing, both lung and kidney tissues were stained with hematoxylin and eosin dyes for morphological examination. Sections were immersed in Harris Hematoxylin (Fisher 67-650-01) for 1 minute, rinsed 2 times in distilled water, differentiated in 0.1% acid alcohol solution for 4 seconds and rinsed in tap water for 15 minutes. Sections were then immersed in eosin for 1 minute, briefly rinsed in tap water, then dehydrated and mounted with Cytoseal Mounting Media (Fisher 23-244-256).

### Immunohistochemistry

After 3 to 6 months, mice were perfused transcardially with PBS, brains were removed and underwent overnight fixation in 70% ethanol in 150 mM NaCl, pH 7.4. After perfusion and fixation, brains were processed into paraffin via sequential dehydration and perfusion with paraffin under vacuum (70% ethanol for 2 hours, 80% ethanol for 1 hour, 95% ethanol for 1 hour, 95% ethanol for 2 hours, 3 times 100% ethanol for 2 hours, xylene for 30 minutes, xylene for 1 hour, xylene for 1.5 hours, 3 times paraffin for 1 hour at 60°C). Brains were then embedded in paraffin blocks, cut into 6 µm sections and mounted on glass slides. Slides were then stained using standard immunohistochemistry as described below. Slides were de-paraffinized with 2 sequential 5-minute washes in xylenes, followed by 1-minute washes in a descending series of ethanols: 100%, 100%, 95%, 80%, 70%. Slides were then incubated in deionized water for one minute prior to microwave antigen retrieval in the BioGenex EZ-Retriever System. Slides were incubated in antigen unmasking solution (Vector Laboratories; Cat# H-3300) and microwaved for 15 minutes at 95°C. Slides were allowed to cool for 20 minutes at room temperature and washed in running tap water for 10 minutes. Slides were incubated in 7.5% hydrogen peroxide in water to quench endogenous peroxidase activity. Slides were washed for 10 minutes in running tap water, 5 minutes in 0.1 M Tris (diluted from 0.5 M Tris made from Tris base and concentrated hydrochloric acid to pH 7.6), then blocked in 0.1 M Tris/2% fetal bovine serum (FBS) for 15 minutes or more. Slides were incubated in primary antibody in 0.1 M Tris/2% FBS in a humidified chamber overnight at 4°C.

To stain pneumocytes, anti-prosurfactant protein C (proSP-C, Millipore Sigma AB3786, RRID: AB_91588) was used at 1:4,000 with microwave antigen retrieval as described above. Primary antibody was rinsed off with 0.1 M tris for 5 minutes, then incubated with goat anti-rabbit (Vector BA1000, RRID: AB_2313606) or horse anti-mouse (Vector BA2000, RRID: AB_2313581) biotinylated IgG in 0.1 M tris/2% FBS 1:1000 for 1 hour. Biotinylated antibody was rinsed off with 0.1 M tris for 5 minutes, then incubated with avidin-biotin solution (Vector PK-6100, RRID: AB_2336819) for 1 hour. Slides were then rinsed for 5 minutes with 0.1 M tris, then developed with ImmPACT DAB peroxidase substrate (Vector SK-4105, RRID: AB_2336520) and counterstained briefly with Harris Hematoxylin (Fisher 67-650-01). Lung ProsP-C data was grouped via object area (μm^2^) and compared with their relative frequencies. To identify statistical differences, this same data was analyzed via beta regression and shifts in the resulting distributions were evaluated using a Kolmogorov-Smirnov test.

For phosphorylated tau, pS202/T205 tau (AT8, ThermoFisher Cat#MN1020, RRID:AB_223647) was used at 1:10,000. Primary antibody was rinsed off with 0.1 M Tris and slides were incubated in 0.1 M Tris for 5 minutes. Primary antibody was detected using the BioGenex Polymer detection kit (Cat #QD440-XAK) per manufacturers protocol as outlined below. Slides were incubated in 50% Enhancer solution in 0.1 M Tris/2% FBS for 20 minutes. Enhancer was rinsed off with 0.1 M Tris, incubated in 0.1 M Tris for 5 minutes, and incubated in 0.1 M Tris/2% FBS for 5 minutes. Slides were then incubated in 50% Poly-HRP in 0.1 M Tris/2% FBS for 30 minutes. Poly-HRP was rinsed off with 0.1 M Tris, slides were then incubated for 5 minutes with 0.1 M Tris, then developed with ImmPACT DAB peroxidase substrate (Vector SK-4105) for 10 minutes. DAB was rinsed off with 0.1 M Tris and incubated in distilled water for 5 minutes. Slides were then counterstained briefly with Harris hematoxylin (ThermoFisher, Cat# 6765001).

Slides were washed in running tap water for 5 minutes, dehydrated in ascending ethanol for 1 minute each (70%, 80%, 95%, 100%, 100%), then washed twice in xylenes for 5 minutes and coverslipped in Cytoseal Mounting Media (Fisher, Cat# 23-244-256). Slides were scanned into digital format on an Aperio AT2 microscope using a 20x objective (0.75 NA) into ScanScope virtual slide (.svs) files. Digitized slides were then used for quantitative pathology.

### Measurement of pS935 and total LRRK2 in kidney lysates

Kidney lysates were prepared in 10 µL/mg Meso Scale Discovery (MSD) lysis buffer (R60TX-2) supplemented with 1x HALT phosphatase and protease inhibitor (Thermo Fisher, 78442). Tissue was placed in a 2 mL round bottom tube (Eppendorf) with a steal bead (Biospec Products, 6.35mm, 11079635C) and homogenized for 1.5 minutes at 30% amplitude in a Qiagen TissueLyser at 40°C. Lysates were centrifuged at 13,000 x *g* for 20 minutes and supernatants were collected for LRRK2 analysis. Total protein was determined with a Micro BCA Protein Assay per kit protocol (Thermo Scientific, 23235). LRRK2 pS935 and total LRRK2 protein levels were quantified from lysates utilizing respective R-PLEX MSD Antibody Sets (LRRK2pS935, F211Q-3; total LRRK2, F211P-3). All incubation steps were performed on a shaker. Streptavidin coated MSD plates (L45SA-2) were coated overnight at 4°C in biotinylated anti-LRRK2pS935 or anti-LRRK2 diluted 1:5 in Diluent 100 (R50AA-3). Plates were washed three times in 1x MSD wash buffer (R61AA-1). The standard curve and samples were applied to the plate and incubated for 1 hour at room temperature. Plates were washed as previously described. SULFO-Tag anti-LRRK2 pS935 or anti-LRRK2 antibodies were diluted 1:100 in Diluent 100, applied to plate and incubated for 1 hour at room temperature. Plates were washed three times, MSD Gold Read Buffer (R92TG-2) was applied, and plates were immediately read on an MSD plate reader.

### Assessment of plasma levels of MLi-2

Plasma was diluted 1:3 in acetonitrile, vortexed for two minutes, and centrifuged at 4,000 RPM for 10 minutes at 40°C. Supernatants were collected and assayed for compound levels using liquid chromatography and tandem mass spectrometry analyses as previously described (Fell et al., 2015).

### Quantitative pathology

All section selection, registration, and quantification were done blinded to treatment. To quantify type II pneumocyte size, we implemented an object classifier in HALO software with a minimum optical density of 0.225. The classifier was able to detect type II pneumocytes well, so these were assayed in the middle of a lung lobe cross-section. Individual object (cell) sizes were measured and the overall distribution of sizes was calculated to determine if there was an overall shift in the size of these cells.

To quantify tau pathology and register brain sections to the Allen Brain Atlas CCFv3, we used an adapted version of the QUINT workflow^27^. Our workflow utilizes a suite of open access programs for segmentation and spatial registration of histological images of the mouse brain—QuPath^42^ for image segmentation, QuickNII^28^ and VisuAlign for image registration, Qmask for hemispheric differentiation and Nutil^29^ for a combination of segmentation and registration and subsequent quantification. Twelve coronal sections per animal were quantified which contain the majority of regions with pathology. To segment pS202/T205 tau, individual sections were segmented using the pixel classification feature of QuPath. An optical density threshold was set at 0.03 to quantify pathology. All pixels above this threshold were quantified, with obvious artifacts removed. The resulting segmentations were exported at full resolution. Down-sampled brain sections were imported into QuickNII and aligned in 3-dimensional space to the 2017 Allen mouse brain atlas CCFv3. Sections were then imported into VisuAlign where anchor points were generated in the atlas and moved to the corresponding location on the section of interest via non-linear warp transformation. In addition, from the spatial coordinate information derived from QuickNII, the masking tool Qmask was used to generate masks over each brain hemisphere so ipsilateral and contralateral brain regions could be analyzed independently. We then used the quantifier feature in Nutil to align the segmentation from QuPath, the registration from VisuAlign, and the hemispheric masks from Qmask to generate percentage area occupied measures for each brain region. The resulting outputs include pixel quantification with spatial location in the brain mapped to 665 brain regions. The resulting output of each of the twelve coronal sections, were then averaged across brain regions to produce single percent area occupied values for each region.

### Computational models of tau pathology spread

Data were modeled under the assumption that tau proteins spread in both anterograde and retrograde directions along the brain’s structural connectome. Structural data were obtained from anterograde viral tract-tracing experiments^43^. The high-resolution structural connectome was constructed by estimating the projection strength between each pair of 100 µm-wide voxels within the Allen Institute CCFv3 whole-brain parcellation^44^. Connections were normalized by dividing the edge strength between each source and target region by the size of the source region^44^.

The equation used to model bidirectional pathology spread is as follows:

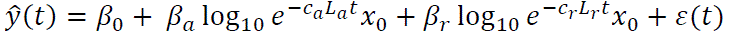

where *b*_0_ is an intercept, *b_a_* is a weight for the importance of anterograde spread, *b_r_* is a weight for the importance of retrograde spread, *c_a_* and *c_r_* are time constants representing the global speed of anterograde and retrograde spread, respectively, *L_a_* and *L_r_* represent the out-degree graph Laplacian for each direction, *t* is time, *x*_0_ is a vector containing ones in the injection site regions (DG, CA1, CA3, VISam, and RSPagl) and zeros elsewhere, and ε is an error term^18^. The “optim” function in R was used to identify the combination of anterograde and retrograde time constants that yielded the strongest correlation between actual and predicted pathology for each group.

To compare model parameters across treatment groups, data from mice in each group were resampled 500 times and each resampled dataset was fit using the bidirectional spread model.

### Model validation

We used previously-described approaches^18^ to evaluate the specificity and performance of our model. To assess the dependence of the model fit on the spatial location of the seed sites, we randomly selected 500 sets of seed regions which had the same average distance from one another as the actual seed sites (1.96 ± 0.196 mm). The bidirectional model described above was then fit for each set of random seed sites, such that *x*_0_ contained ones in the randomly-selected seed regions and zeros elsewhere.

To evaluate the performance of the bidirectional spread model, we compared fits obtained from models accounting for spread in only one direction (anterograde or retrograde) or based solely on the distance between regions. For each model type, data from mice of both genotypes and treatment groups were pooled and randomly resampled to obtain *y_train_*(*t*) and *y_test_*(*t*) for each time point. The model fit was performed on *y_train_*(*t*) and the fit with *y_test_*(*t*) was then evaluated. This process was repeated 500 times for each model type to obtain distributions of out-of-sample fits.

### Statistical analysis

The number of animals analyzed in each experiment, the statistical analysis performed, as well as the p-values for all results <0.05 are reported in the figure legends. *In vivo* pathological spread data was analyzed and all computations were performed in R (https://www.R-project.org/)^45^ as described.

For analyses assessing differences in percent area occupied, data were stratified based on brain hemisphere (ipsilateral/contralateral), brain region, MPI, and either genotype or treatment depending on the specific hypothesis test (i.e. stratified on treatment if testing genotype differences and vice-versa). Regions with zero variance across all treatments and genotypes were filtered out. To maintain flexibility and avoid reliance on strong parametric assumptions, robust linear regressions via the *MASS* package, with the ranked percent area occupied as the outcome and either genotype or treatment as the explanatory variable, were used (https://cran.r-project.org/web/packages/MASS/index.html). All models were adjusted for sex and daughter regions when the number of daughter regions was 2 or greater. Effect sizes, including confidence intervals were estimated using the R package *emmeans* (R package version 1.8.3, URL: https://CRAN.R-project.org/package=emmeans). Significance was determined using second generation p-values based on a null interval of +/-5% difference 95% confidence intervals ^46^. Only second-generation p-values equal to 0 were considered significant.

## Supplemental Information for

**Supplementary Table 1.** Characterization of PHF preparations from AD brains. Three cases were extracted for this study. Information, including the post-mortem interval (PMI) for these cases is displayed in the table, in addition to protein concentrations used to evaluate the amount of protein to inject.

| Case | Age | Sex | PMI (hours) | Total protein ($\mu\text{g}/\mu\text{L}$) | Tau ($\mu\text{g}/\mu\text{L}$) | Tau (% protein) | α-Syn ($\mu\text{g}/\text{mL}$) | Aβ 1-40 (ng/mL) | Aβ 1-42 (ng/mL) |
| --- | --- | --- | --- | --- | --- | --- | --- | --- | --- |
| **AD Case 1** | 89 | F | 18 | 7.1 | 0.67 | 9.4 | 0.620 | 10.74 | 102.71 |
| **AD Case 2** | 74 | M | 18 | 6.7 | 0.74 | 11.0 | 0.024 | 3.57 | 57.23 |
| **AD Case 3** | 82 | F | 7 | 4.6 | 0.45 | 9.8 | 0.029 | 16.79 | 131.53 |

**Supplementary Figure 1.**
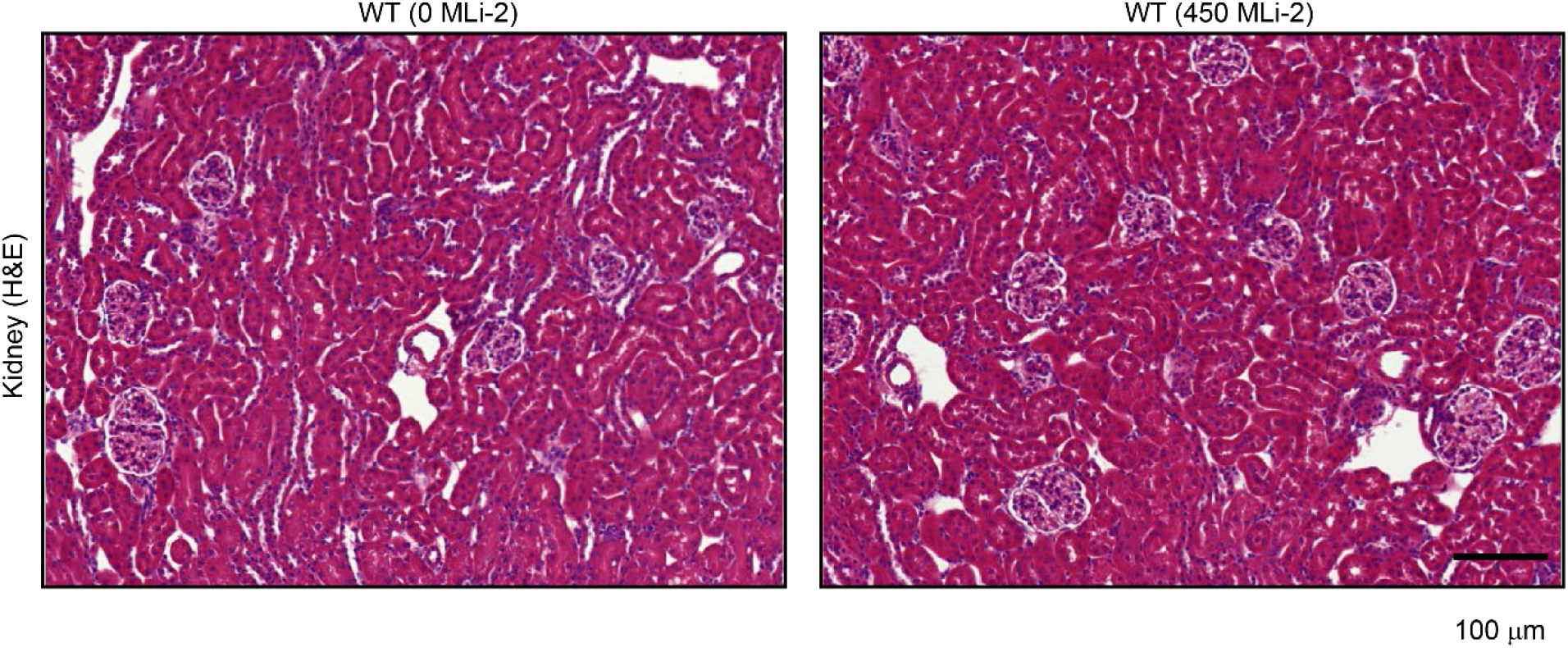
Long-term MLi-2 does not impact gross kidney morphology. All experimental animals had their kidneys removed, sectioned, and stained by hematoxylin and eosin. No gross morphological differences were apparent in any of the cohorts. As an example, kidneys from wildtype mice treated with 0 mg/kg MLi-2 are compared above to wildtype mice treated with 450 mg/kg MLi-2 for 6 months. Scale bar = 100 µm.

**Supplementary Figure 2.**
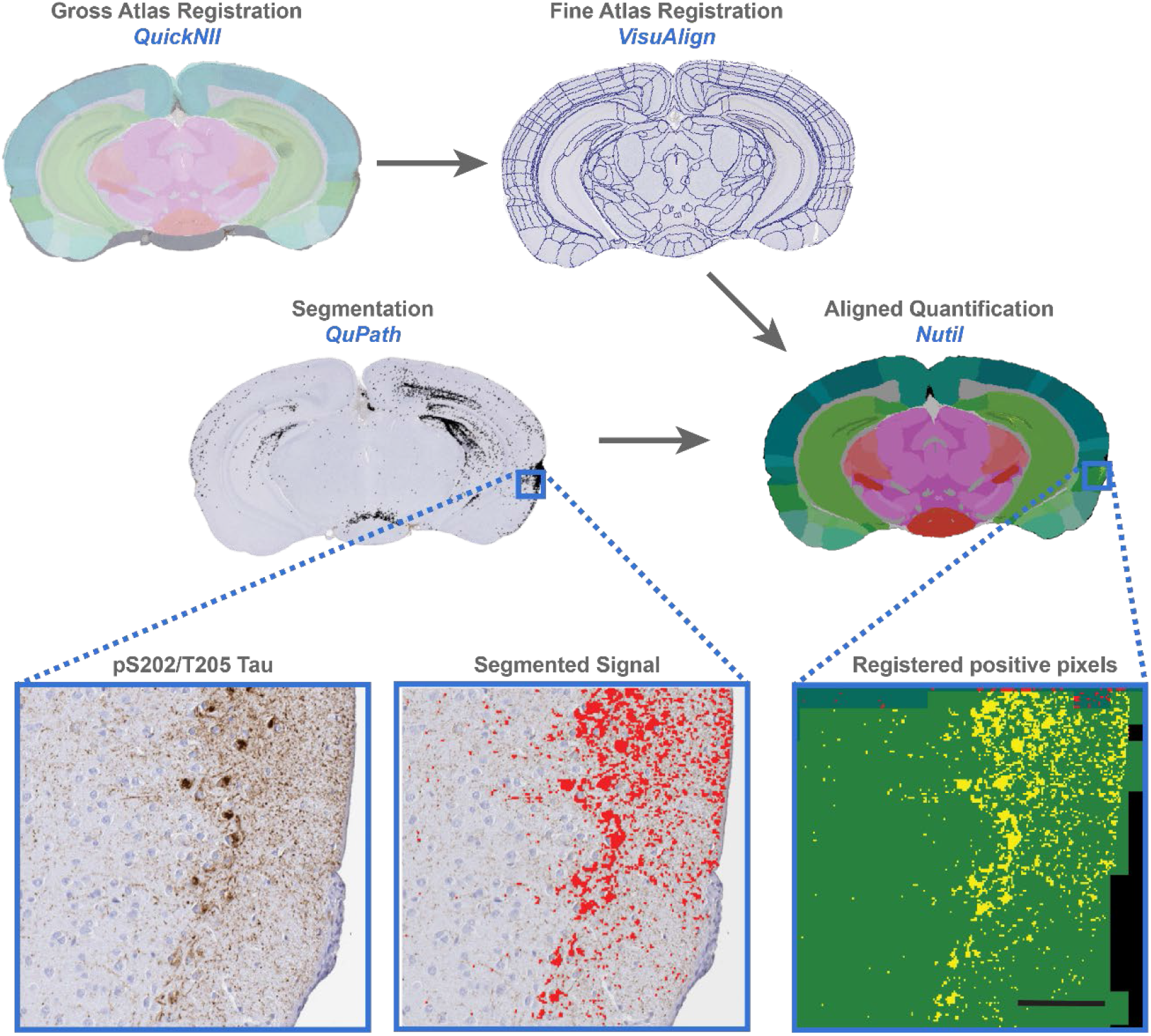
Quantitative pathology workflow. Brains from mice injected with AD PHFs were mounted in paraffin blocks, sectioned and stained for pathological tau (AT8, pS202/T205). Stained tissue was then scanned into a digitized format. Segmentation and registration to the Allen Brain Atlas CCFv3 was performed with a modified version of the QUINT workflow. Pathology was segmented in QuPath based on pixel intensity. A parallel image was registered to the Allen Brain Atlas using QuickNII software for gross registration and VisuAlign for fine registration, including non-linear warp transformation. Finally, segmentation and registration data were merged in Nutil to provide outputs of percentage of areas occupied for each brain region. Representative pathology segmentation and registration are shown in enlarged images for the entorhinal region of this brain. Scale bar = 100 µm.

**Supplementary Figure 3.**
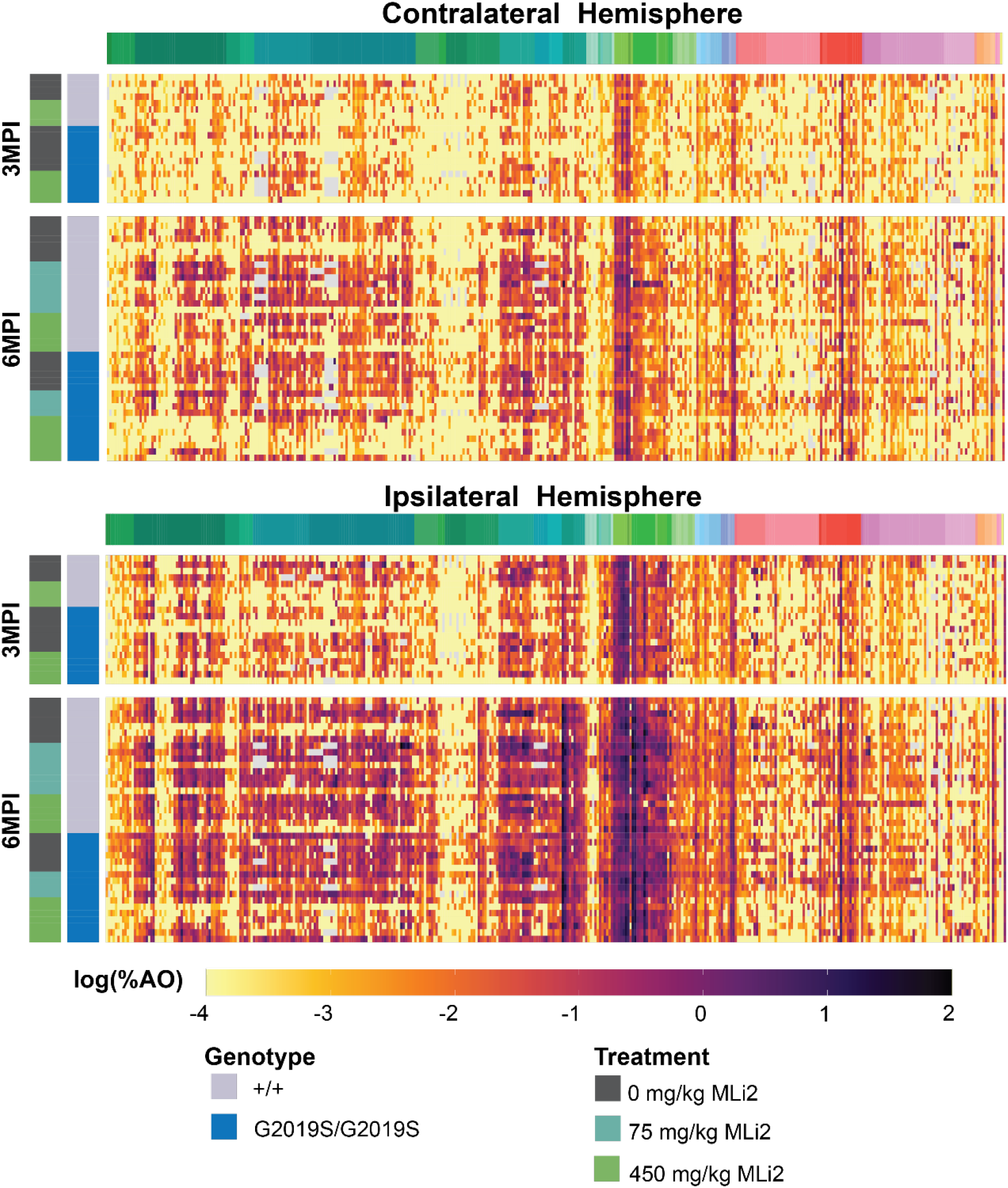
Quantitative pathology analysis from all mice. Log percentage area occupied is plotted for all measured regions and mice, with the treatment and genotypes noted on the x-axis, and the brain region denoted on the y-axis by Allen Brain Atlas region coloration.

**Supplementary Figure 4.**
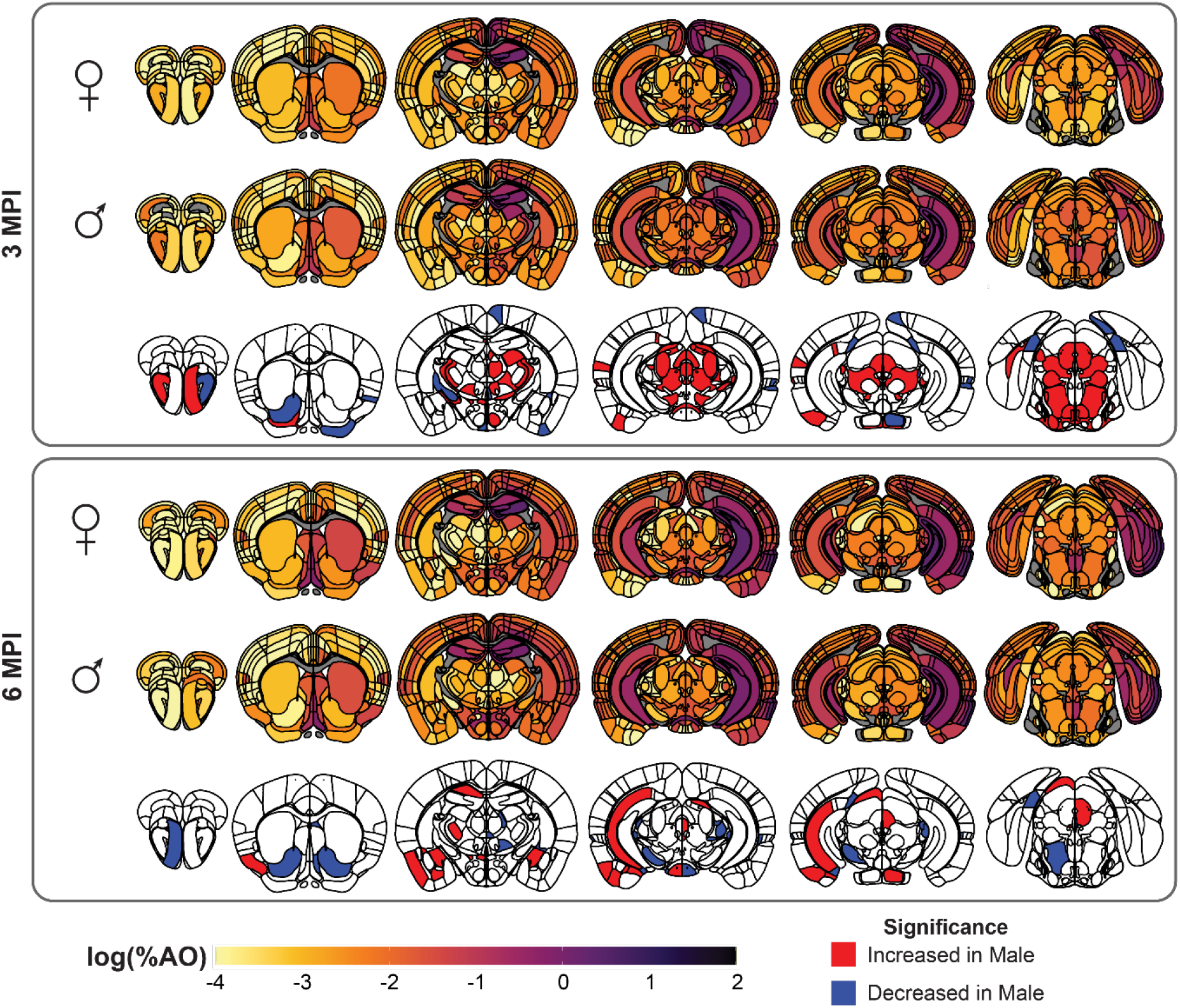
Sex differences in tau pathology in wildtype mice. Anatomical heatmaps of average log percentage area occupied with pS202/T205 tau is plotted for male and female mice at 3 and 6 months post-injection and second-generation *p*-values of regional statistical significance of male compared to female mice (δp=0). While the majority of cortical regions show no different between males and females, several subcortical regions show differences. Most notable are the increase in thalamic and mesencephalic regions and CA1 region of the contralateral hippocampus in male mice.

**Supplementary Figure 5.**
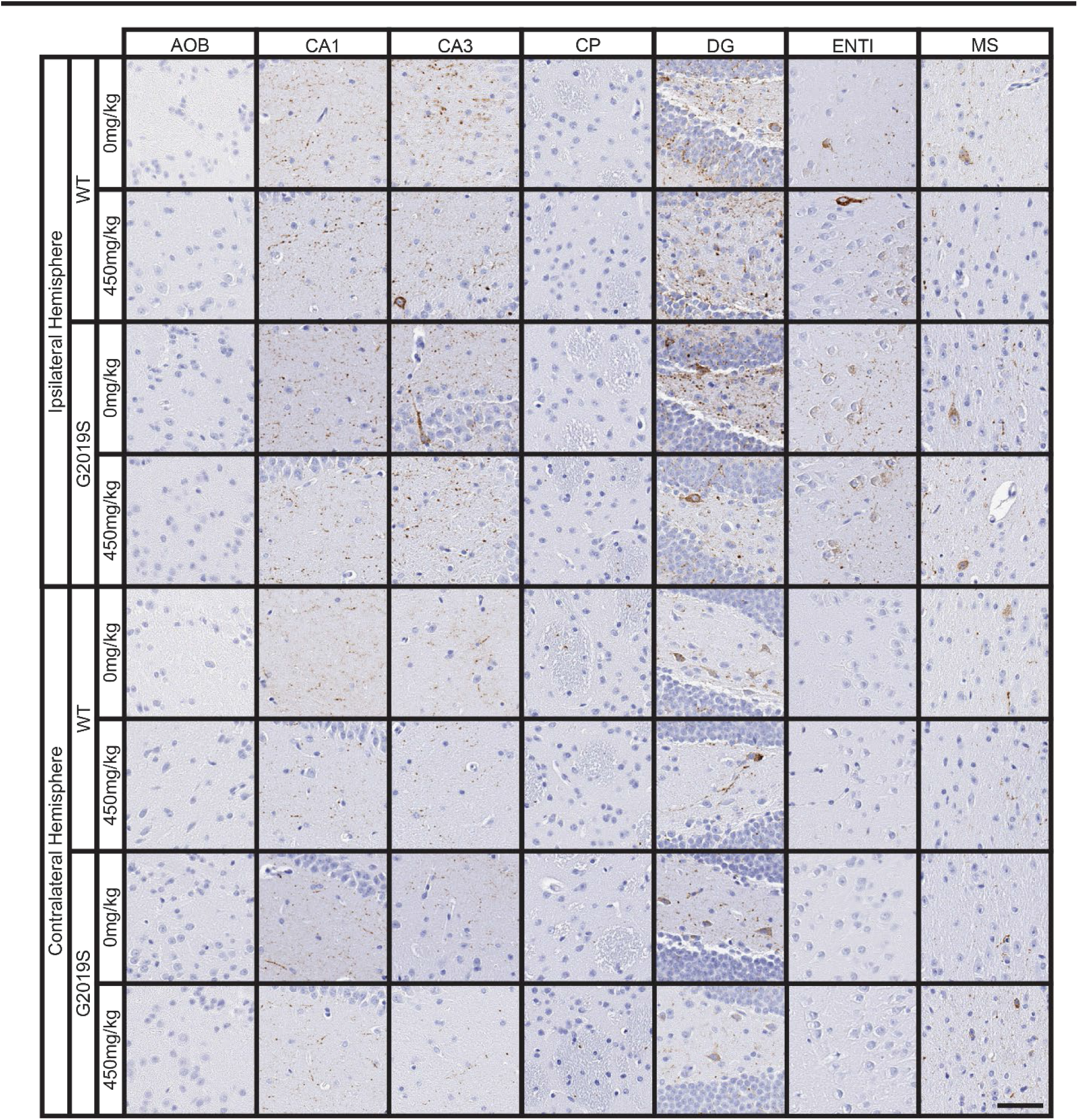
Representative staining from 3 MPI mice. Representative images of selected regional tau pathology from 3 MPI mice. AOB: accessory olfactory bulb, CA1: field CA1 of hippocampus, CA3: field CA3 of hippocampus, CP: caudoputamen, DG: dentate gyrus, ENTl: entorhinal area, lateral part, MS: medial septal nucleus. Scale bar = 50 μm.

**Supplementary Figure 6.**
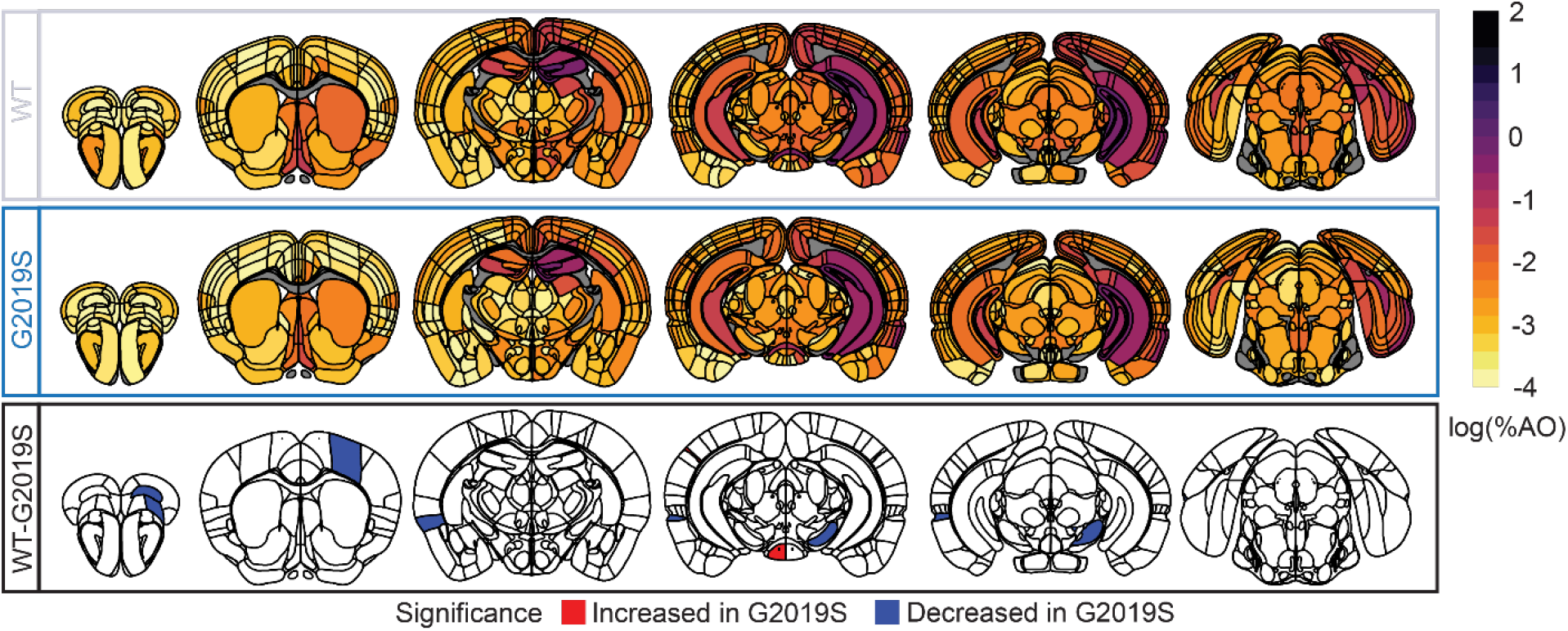
Wildtype compared to LRRK2^G2019S^ mice at 3 MPI. Anatomic heatmaps of mean regional tau pathology shown as log (% area occupied) at 3 MPI and second-generation *p*-values of regional statistical significance of wildtype mice compared to G2019S mice (δp=0). Tau pathology was not quantified in white matter regions, so they are plotted as gray. There were minimal differences between genotypes at this timepoint.

**Supplementary Figure 7.**
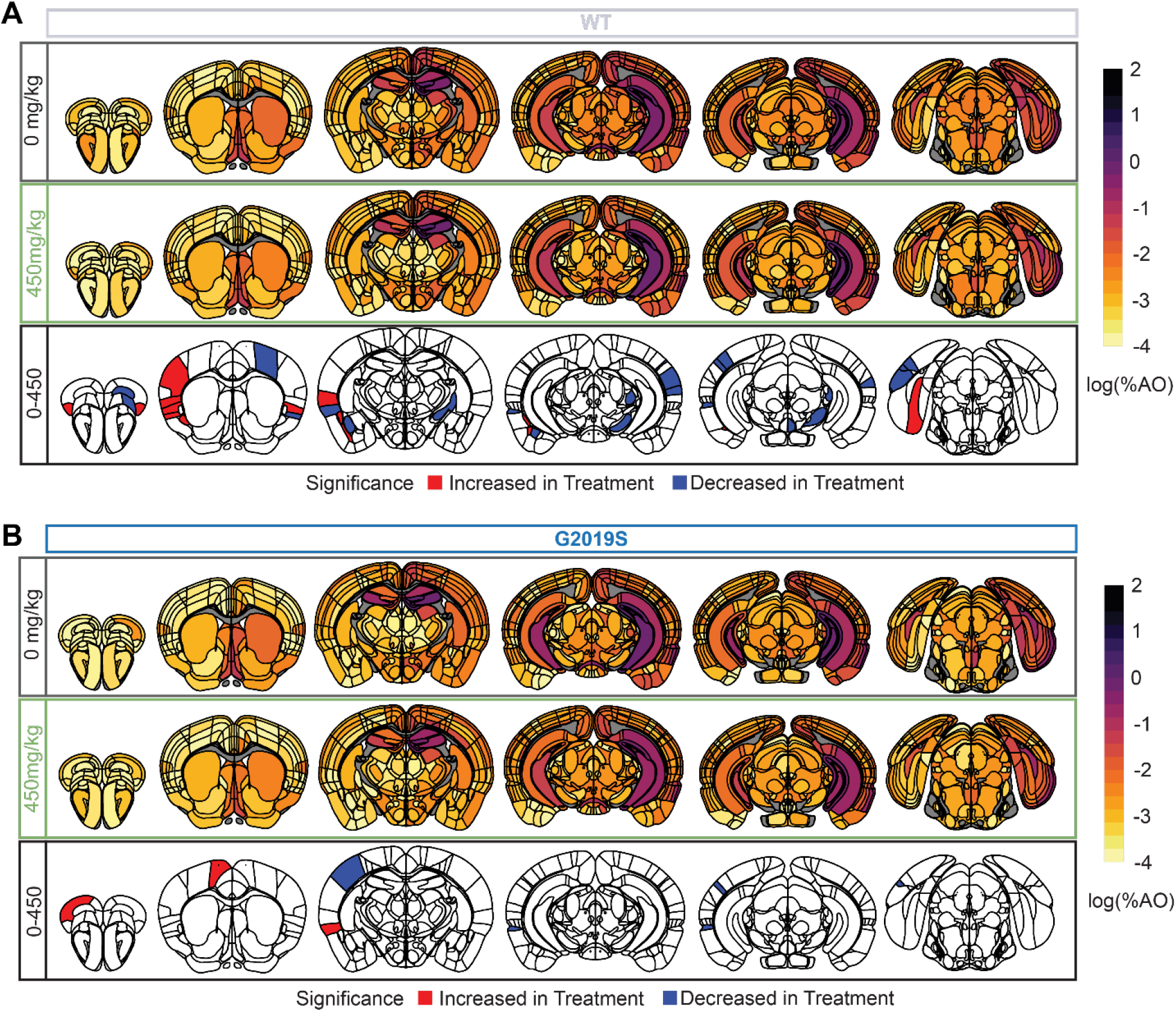
Tau pathology compared by treatment group at 3 MPI. (**A**) Anatomic heatmaps of mean regional tau pathology shown as log (% area occupied) at 3 MPI and second-generation *p*-values of regional statistical significance of wildtype mice treated with 0 mg/kg MLi-2 compared to 450 mg/kg MLi-2 (δp=0). (**B**) Anatomic heatmaps of mean regional tau pathology shown as log (% area occupied) at 3 MPI and second-generation *p*-values of regional statistical significance of LRRK2^G2019S^ mice treated with 0 mg/kg MLi-2 compared to 450 mg/kg MLi-2 (δp=0). There were minimal differences associated with treatment at this timepoint.

**Supplementary Figure 8.**
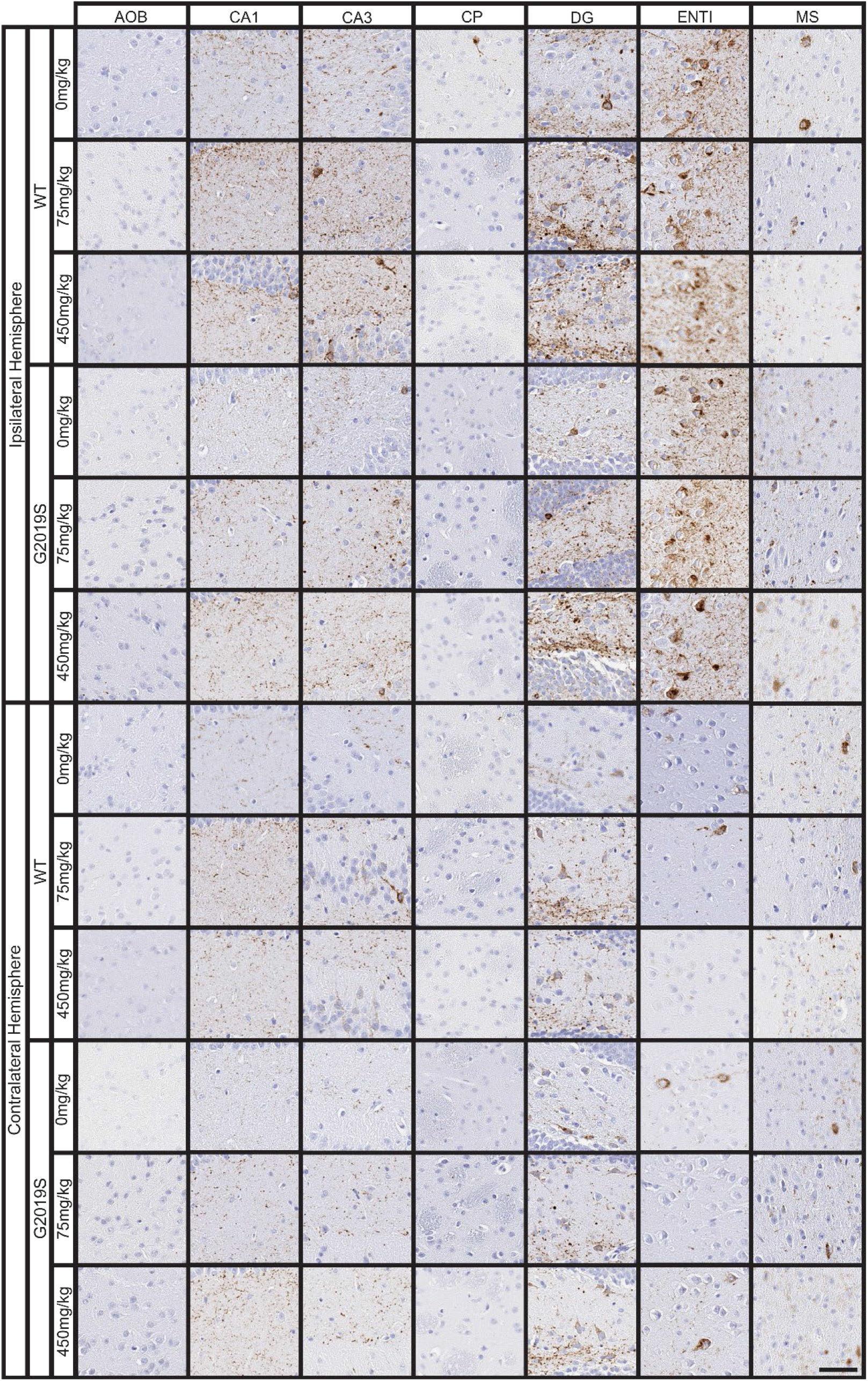
Representative staining from 6 MPI mice. Representative images of selected regional tau pathology (AT8, pS202/T205) from 6 MPI mice. Scale bar = 50 μm.

**Supplementary Figure 9.**
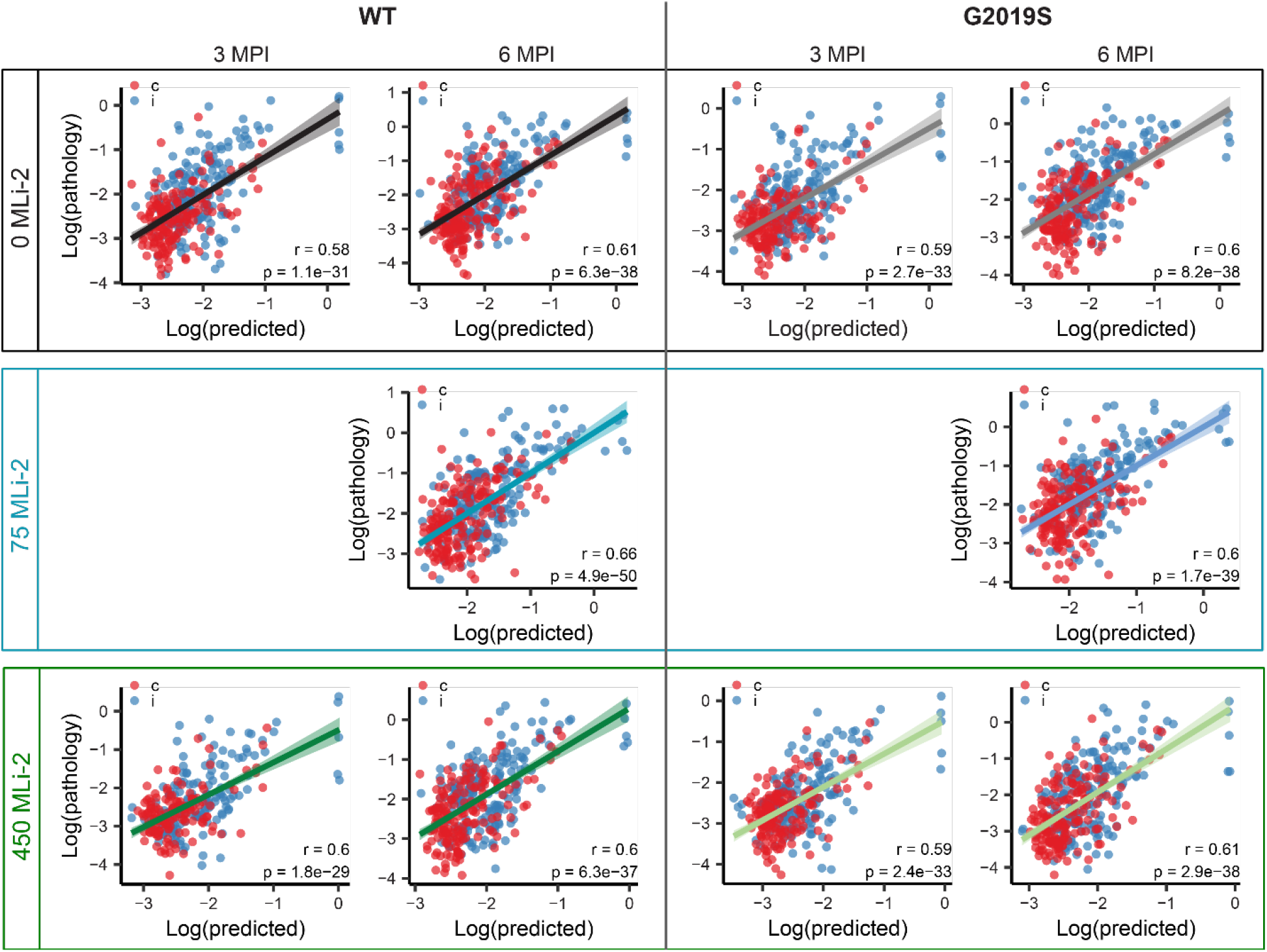
Examination of linear diffusion fits by hemisphere. Predictions of log tau pathology from linear diffusion models based on bidirectional (anterograde and retrograde) anatomical connections. Solid lines represent the line of best fit, and shading represents 95% confidence intervals. This is the same data as plotted in Fig. 6A, except ipsilateral (blue) and contralateral (red) brain regions are colored differently.

## REFERENCES

1 Goedert, M., Spillantini, M. G., Del Tredici, K. & Braak, H. 100 years of Lewy pathology. Nature reviews. Neurology 9, 13–24, doi:10.1038/nrneurol.2012.242 (2013).

2 Healy, D. G. et al. Phenotype, genotype, and worldwide genetic penetrance of LRRK2-associated Parkinson’s disease: a case-control study. The Lancet. Neurology 7, 583–590, doi:10.1016/S1474-4422(08)70117-0 (2008).

3 Rocha, E. M., Keeney, M. T., Di Maio, R., De Miranda, B. R. & Greenamyre, J. T. LRRK2 and idiopathic Parkinson’s disease. Trends in neurosciences 45, 224–236, doi:10.1016/j.tins.2021.12.002 (2022).

4 Lee, A. J. et al. Penetrance estimate of LRRK2 p.G2019S mutation in individuals of non-Ashkenazi Jewish ancestry. Movement disorders: official journal of the Movement Disorder Society 32, 1432–1438, doi:10.1002/mds.27059 (2017).

5 Azeggagh, S. & Berwick, D. C. The development of inhibitors of leucine-rich repeat kinase 2 (LRRK2) as a therapeutic strategy for Parkinson’s disease: the current state of play. British journal of pharmacology 179, 1478–1495, doi:10.1111/bph.15575 (2022).

6 Di Maio, R. et al. LRRK2 activation in idiopathic Parkinson’s disease. Science translational medicine 10, doi:10.1126/scitranslmed.aar5429 (2018).

7 Rodrigues e Silva, A. M., et al. Who was the man who discovered the “Lewy bodies”? Mov Disord 25, 1765–1773, doi:10.1002/mds.22956 (2010).

8 Irwin, D. J. et al. Neuropathological and genetic correlates of survival and dementia onset in synucleinopathies: a retrospective analysis. The Lancet. Neurology 16, 55–65, doi:10.1016/S1474-4422(16)30291-5 (2017).

9 Poulopoulos, M., Levy, O. A. & Alcalay, R. N. The neuropathology of genetic Parkinson’s disease. Movement disorders: official journal of the Movement Disorder Society 27, 831–842, doi:10.1002/mds.24962 (2012).

10 Kalia, L. V. et al. Clinical correlations with Lewy body pathology in LRRK2-related Parkinson disease. JAMA neurology 72, 100–105, doi:10.1001/jamaneurol.2014.2704 (2015).

11 Henderson, M. X., Sengupta, M., Trojanowski, J. Q. & Lee, V. M. Y. Alzheimer’s disease tau is a prominent pathology in LRRK2 Parkinson’s disease. Acta neuropathologica communications 7, 183, doi:10.1186/s40478-019-0836-x (2019).

12 Blauwendraat, C. et al. Genetic analysis of neurodegenerative diseases in a pathology cohort. Neurobiol Aging 76, 214 e211–214 e219, doi:10.1016/j.neurobiolaging.2018.11.007 (2019).

13 Ross, O. A. et al. Lrrk2 R1441 substitution and progressive supranuclear palsy. Neuropathology and applied neurobiology 32, 23–25, doi:10.1111/j.1365-2990.2006.00693.x (2006).

14 Sanchez-Contreras, M. et al. Study of LRRK2 variation in tauopathy: Progressive supranuclear palsy and corticobasal degeneration. Movement disorders: official journal of the Movement Disorder Society 32, 115–123, doi:10.1002/mds.26815 (2017).

15 Bieri, G. et al. LRRK2 modifies alpha-syn pathology and spread in mouse models and human neurons. Acta neuropathologica 137, 961–980, doi:10.1007/s00401-019-01995-0 (2019).

16 Henderson, M. X. et al. Spread of alpha-synuclein pathology through the brain connectome is modulated by selective vulnerability and predicted by network analysis. Nat Neurosci 22, 1248–1257, doi:10.1038/s41593-019-0457-5 (2019).

17 Volpicelli-Daley, L. A. et al. G2019S-LRRK2 Expression Augments alpha-Synuclein Sequestration into Inclusions in Neurons. J Neurosci 36, 7415–7427, doi:10.1523/JNEUROSCI.3642-15.2016 (2016).

18 Cornblath, E. J. et al. Computational modeling of tau pathology spread reveals patterns of regional vulnerability and the impact of a genetic risk factor. Sci Adv 7, doi:10.1126/sciadv.abg6677 (2021).

19 Nguyen, A. P. T. et al. 369 G2019S LRRK2 enhances the neuronal transmission of tau in the mouse brain. Hum Mol Genet 27, 120–134, doi:10.1093/hmg/ddx389 (2018).

20 Fell, M. J. et al. MLi-2, a Potent, Selective, and Centrally Active Compound for Exploring the Therapeutic Potential and Safety of LRRK2 Kinase Inhibition. The Journal of pharmacology and experimental therapeutics 355, 397–409, doi:10.1124/jpet.115.227587 (2015).

21 Matikainen-Ankney, B. A. et al. Altered Development of Synapse Structure and Function in Striatum Caused by Parkinson’s Disease-Linked LRRK2-G2019S Mutation. J Neurosci 36, 7128–7141, doi:10.1523/JNEUROSCI.3314-15.2016 (2016).

22 Guo, J. L. et al. Unique pathological tau conformers from Alzheimer’s brains transmit tau pathology in nontransgenic mice. The Journal of experimental medicine 213, 2635–2654, doi:10.1084/jem.20160833 (2016).

23 Bryce, D. K. et al. Characterization of the Onset, Progression, and Reversibility of Morphological Changes in Mouse Lung after Pharmacological Inhibition of Leucine-Rich Kinase 2 Kinase Activity. The Journal of pharmacology and experimental therapeutics 377, 11–19, doi:10.1124/jpet.120.000217 (2021).

24 Fuji, R. N. et al. Effect of selective LRRK2 kinase inhibition on nonhuman primate lung. Science translational medicine 7, 273ra215, doi:10.1126/scitranslmed.aaa3634 (2015).

25 Andersen, M. A. et al. PFE-360-induced LRRK2 inhibition induces reversible, non-adverse renal changes in rats. Toxicology 395, 15–22, doi:10.1016/j.tox.2018.01.003 (2018).

26 Henderson, M. X. et al. LRRK2 inhibition does not impart protection from alpha-synuclein pathology and neuron death in non-transgenic mice. Acta neuropathologica communications 7, 28, doi:10.1186/s40478-019-0679-5 (2019).

27 Yates, S. C. et al. QUINT: Workflow for Quantification and Spatial Analysis of Features in Histological Images From Rodent Brain. Frontiers in neuroinformatics 13, 75, doi:10.3389/fninf.2019.00075 (2019).

28 Puchades, M. A., Csucs, G., Ledergerber, D., Leergaard, T. B. & Bjaalie, J. G. Spatial registration of serial microscopic brain images to three-dimensional reference atlases with the QuickNII tool. PloS one 14, e0216796, doi:10.1371/journal.pone.0216796 (2019).

29 Groeneboom, N. E., Yates, S. C., Puchades, M. A. & Bjaalie, J. G. Nutil: A Pre– and Post-processing Toolbox for Histological Rodent Brain Section Images. Frontiers in neuroinformatics 14, 37, doi:10.3389/fninf.2020.00037 (2020).

30 Henderson, M. X. et al. Glucocerebrosidase Activity Modulates Neuronal Susceptibility to Pathological alpha-Synuclein Insult. Neuron 105, 822–836 e827, doi:10.1016/j.neuron.2019.12.004 (2020).

31 Tolosa, E., Vila, M., Klein, C. & Rascol, O. LRRK2 in Parkinson disease: challenges of clinical trials. Nature reviews. Neurology 16, 97–107, doi:10.1038/s41582-019-0301-2 (2020).

32 Baptista, M. A. S. et al. LRRK2 inhibitors induce reversible changes in nonhuman primate lungs without measurable pulmonary deficits. Science translational medicine 12, doi:10.1126/scitranslmed.aav0820 (2020).

33 Jennings, D. et al. Preclinical and clinical evaluation of the LRRK2 inhibitor DNL201 for Parkinson’s disease. Science translational medicine 14, eabj2658, doi:10.1126/scitranslmed.abj2658 (2022).

34 Dues, D. J. et al. Formation of templated inclusions in a forebrain alpha-synuclein mouse model is independent of LRRK2. bioRxiv, doi:10.1101/2023.08.19.553965 (2023).

35 Beccano-Kelly, D. A. et al. Synaptic function is modulated by LRRK2 and glutamate release is increased in cortical neurons of G2019S LRRK2 knock-in mice. Frontiers in cellular neuroscience 8, 301, doi:10.3389/fncel.2014.00301 (2014).

36 Boecker, C. A., Goldsmith, J., Dou, D., Cajka, G. G. & Holzbaur, E. L. F. Increased LRRK2 kinase activity alters neuronal autophagy by disrupting the axonal transport of autophagosomes. Current biology: CB 31, 2140–2154 e2146, doi:10.1016/j.cub.2021.02.061 (2021).

37 Vogel, J. W. et al. Connectome-based modelling of neurodegenerative diseases: towards precision medicine and mechanistic insight. Nature reviews. Neuroscience 24, 620–639, doi:10.1038/s41583-023-00731-8 (2023).

38 van Dyck, C. H. et al. Lecanemab in Early Alzheimer’s Disease. The New England journal of medicine 388, 9–21, doi:10.1056/NEJMoa2212948 (2023).

39 Scholl, M. et al. PET Imaging of Tau Deposition in the Aging Human Brain. Neuron 89, 971–982, doi:10.1016/j.neuron.2016.01.028 (2016).

40 Schwarz, A. J. et al. Regional profiles of the candidate tau PET ligand 18F-AV-1451 recapitulate key features of Braak histopathological stages. Brain: a journal of neurology 139, 1539–1550, doi:10.1093/brain/aww023 (2016).

41 Irwin, D. J. et al. Neuropathologic substrates of Parkinson disease dementia. Ann Neurol 72, 587–598, doi:10.1002/ana.23659 (2012).

42 Bankhead, P. et al. QuPath: Open source software for digital pathology image analysis. Sci Rep 7, 16878, doi:10.1038/s41598-017-17204-5 (2017).

43 Oh, S. W. et al. A mesoscale connectome of the mouse brain. Nature 508, 207–214, doi:10.1038/nature13186 (2014).

44 Knox, J. E. et al. High-resolution data-driven model of the mouse connectome. Netw Neurosci 3, 217–236, doi:10.1162/netn_a_00066 (2019).

45 R: A Language and Environment for Statistical Computing (2018).

46 Blume, J. D., Greevy, R. A., Welty, V. F., Smith, J. R. & Dupont, W. D. An Introduction to Second-Generation p-Values. The American Statistician 73, 157–167, doi:10.1080/00031305.2018.1537893 (2019).

